# Inducing chondrogenesis in MSC/chondrocyte co-cultures using exogenous TGF-*β*: a mathematical model

**DOI:** 10.1101/141119

**Authors:** M.J. Chen, J.P. Whiteley, C.P. Please, A. Schwab, F. Ehlicke, S.L. Waters, H.M. Byrne

## Abstract

The differentiation of mesenchymal stem cells (MSCs) into chondrocytes (native cartilage cells), or chondrogenesis, is a key step in the tissue engineering of articular cartilage, where the motility and high proliferation rate of MSCs used as seed cells are exploited. Chondrogenesis is regulated by transforming growth factor-beta (TGF-*β*), a short-lived cytokine whose effect is prolonged by storage in the extracellular matrix. Tissue engineering applications require the complete differentiation of an initial population of MSCs, and two common strategies used to achieve this *in vitro* are (1) co-culture the MSCs with chondrocytes, which constitutively produce TGF-*β*; or (2) add exogenous TGF-*β*. To investigate these strategies we develop an ordinary differential equation model of the interactions between TGF-*β*, MSCs and chondrocyte. Here the dynamics of TGF-*β* are much faster than those of the cell processes; this difference in time-scales is exploited to simplify subsequent model analysis. Using our model we demonstrate that under strategy 1 complete chondrogenesis will be induced if the initial proportion of chondrocytes exceeds a critical value. Similarly, under strategy 2 we find that there is a critical concentration of exogenous TGF-*β* above which all MSCs will ultimately differentiate. Finally, we use the model to demonstrate the potential advantages of adopting a hybrid strategy where exogenous TGF-*β* is added to a co-culture of MSCs and chondrocytes, as compared to using either strategy 1 or 2 in isolation.

## 1. Introduction

Articular cartilage is a thin layer of dense tissue found on the ends of bones in synovial joints. It prevents contact between the ends of the bones and acts as a lubricating surface between them [1, 2]. Cartilage is composed of a combination of native cartilage cells (or chondrocytes) and extracellular matrix components, principally collagen and proteoglycans. The zonated structure and arrangement of both the cells and the matrix components endow cartilage with properties that make it particularly well suited to withstanding mechanical stresses in load bearing joints, such as the knee (a schematic cross-section of this zonated structure is shown in Figure 1). Articular cartilage has a low capacity to regenerate due to its avascular structure and the low motility and proliferation rates of the chondrocytes. Thus, any degeneration of cartilage is chronic and can lead to severe conditions, such as osteoarthritis, which typically require some form of surgical intervention. A proposed alternative to current joint replacement approaches involves implanting artificially engineered cartilage to replace the damaged tissue.

**Figure 1:**
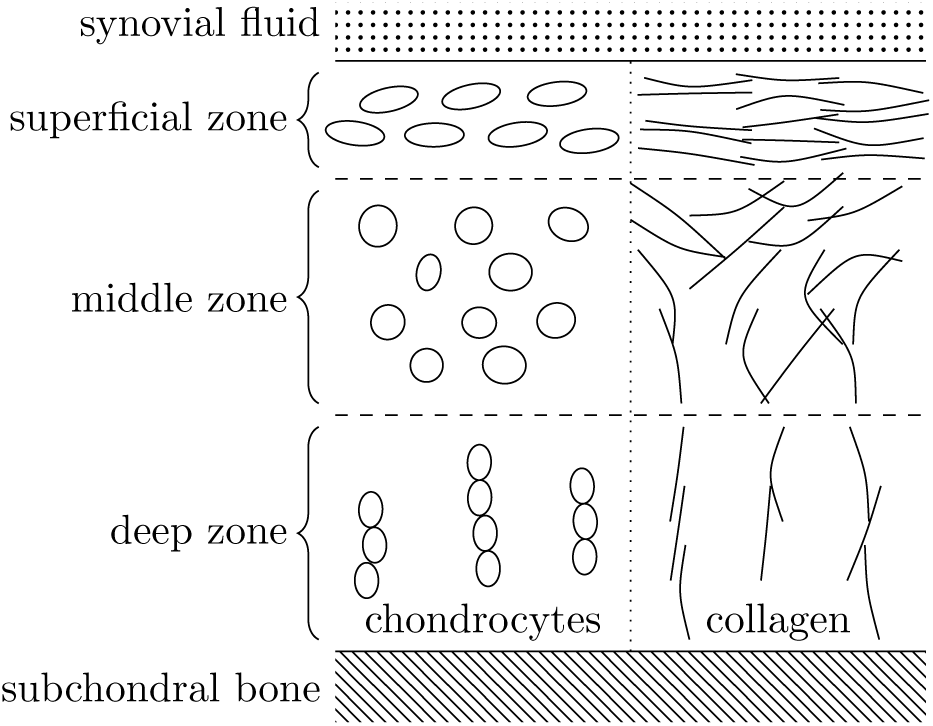
Schematic cartoon of the zonated, depth-dependent structure of articular cartilage. In a synovial joint the cartilage lies between the synovial fluid and the subchondral bone, and is typically described as being divided into three zones. The highest density of both collagen and chondrocytes is in the superficial zone, and here both are aligned in the direction of the surface of the cartilage. In the middle zone the cells are more rounded and the alignment of the collagen is less organised, with the density of both lower than in the superficial zone. In the deep zone the cells are organised in column-like structures, with both these columns and the collagen aligned perpendicular to the subchondral bone [2].

Ideally such an implant should mimic the biomechanical function of the natural tissue and have a similar distribution of cells and matrix components. One promising effort to produce such an implant involves seeding cells in a hydrogel construct reinforced with a lattice of 3D-printed polymer fibres, which is then cultured in a bioreactor under chemical and mechanical stimuli [3]. The seeded cells may be either chondrocytes harvested from natural cartilage, or mesenchymal stem cells (MSCs), which differentiate into chondrocytes in response to externally imposed stimuli, or a combination of both cell types. The relevant matrix components are synthesised and maintained by chondrocytes, so a zonated distribution of these components requires a zonated distribution of the chondrocytes. As chondrocytes have very low motility this distribution may be achieved either by seeding chondrocytes in a zonated fashion, or by exploiting the high motility of the MSCs and stimulating them to move throughout the construct before they differentiate.

A further consideration is the low proliferation rate of chondrocytes which means that many cells must be harvested from other sources to seed the construct; such harvesting is practically difficult to achieve. By contrast, MSCs are highly proliferative, so fewer cells need be harvested; the challenges in their use are to bias their differentiation into chondrocytes (chondrogenesis) and then maintain a stable phenotype [4] (we note that MSCs may also differentiate into bone or muscle cells, for instance). Chemical stimulation can be provided to the seeded cells, for instance in the form of fetal bovine serum (FBS), platelet-derived growth factor (PDGF), or transforming growth factor-*β* (TGF-*β*), to promote proliferation, differentiation and/or chemotactic movement of the MSCs [5, 6, 7]. Chemical stimulation can be combined with mechanical stimulation, for instance by using a piston to apply a load to the hydrogel construct [8], to promote cell differentiation via mechanotransduction.

Identifying appropriate conditions under which to culture artificial constructs to produce a specific end-product is technically challenging. The cells are typically highly sensitive to small changes in the environment, and therefore a small change in the concentration of a particular growth factor may have a large effect. Similarly, for mechanical loading a balance needs to be struck between applying enough loading to induce mechanotransductive effects, but not too much that the structure of the hydrogel is compromised. Additionally, in the context of tissue engineering differentiated cells may dedifferentiate, and so may require regular stimulation to prevent their loss of phenotype. Producing a desired outcome involves carefully balancing the application of the different stimuli and the responses they induce in the cells. The modelling work of this study aims to address this challenging question. We develop a mathematical model to investigate how to drive a population of MSCs to chondrogenesis with a focus on the response of MSCs to a single growth factor in particular, transforming growth factor *β* (TGF-*β*).

TGF-*β* is particularly important in a number of contexts related to cartilage. *In vivo* it is produced by chondrocytes and stored bound to the ECM. In unbound form it stimulates chondrocytes to synthesise ECM components such as collagen type II and proteoglycans (particularly aggrecan). It drives MSCs to differentiate into chondrocytes. The biochemistry of TGF-*β* has several subtleties: it is secreted in active or latent form as part of a ‘large latent complex’, whose active component is bound to a protein and a peptide and unable to bind to cell receptors until it is released from this complex. This arrangement enables the latent TGF-*β* to bind to and be stored in the ECM, for release only when chondrocyte stimulation is required (in response to ECM damage for instance). TGF-*β* activation typically occurs in response to mechanical or chemical stimulation [9, 10]. Once activated it has a short half-life of only a few minutes as compared to the time-scale of days over which cell differentiation and proliferation occur [11]. Consequently it must bind rapidly to cells after activation to have any effect. For *in vitro* studies, TGF-*β* is replenished at regular intervals to ensure a sustained effect. For tissue engineering applications there are two dominant sources of TGF-*β*: it may be secreted by chondrocytes or it may be added exogenously. The key modelling question that we address in this paper is then to understand how these two sources interact and, in particular, to establish whether exogenous TGF-*β* alone is sufficient to drive stem cell differentiation.

A number of experimental strategies have been developed to drive the chondrogenesis of an *in vitro* population of MSCs. One strategy involves adding exogenous TGF-*β* to the initial population of stem cells [8]. Although TGF-*β* has a short half life, if its initial concentration is high enough it is possible that it will drive some cells to differentiate before it degrades completely. These newly differentiated chondrocytes will constitutively produce more TGF-*β*, and if they do so in a large enough quantity then this may drive further differentiation. Where too few cells are initially differentiated more TGF-*β* may be added to achieve complete differentiation. Mathematical modelling of this excitable system will help to identify conditions under which the system can be driven to constitutively produce enough TGF-*β* to sustain cell differentiation after the initial source of exogenous TGF-*β* has been depleted.

An alternative strategy for obtaining a population of differentiated cells is to co-culture MSCs with some harvested chondrocytes. There is evidence that co-cultures of this type produce cell populations with a more stable phenotype [12]. Under these conditions, the mechanism that induces chondrogenesis is similar to that above, except that the only source of the TGF-*β* is that produced by the chondrocytes. A point of interest here is how few chondrocytes need to be added to induce chondrogenesis of the whole cell population, since chondrocytes are difficult to harvest.

We have performed experiments to investigate this chondrogenesis strategy and an example is shown in Fig. 2. Here, a co-culture of 80% MSCs (at a density of 16 Mio/mL) to 20% chondrocytes (at a density of 4 Mio/mL) yielded comparable levels of chondrogenic markers (shown here are stainings for collagen type II) after 28 days to that of a population of 100% chondrocytes (at a density of 20 Mio/mL). This indicates that the MSCs have differentiated and are producing this particular ECM component at a rate similar to the seeded native cartilage cells. If exogenous TGF-*β* is added to a mixed population of cells then it is possible that even fewer chondrocytes and a lower concentration of exogenous TGF-*β* will be required than if the these strategies are used in isolation [13]. Mathematical models of this scenario can be used to assess the relative importance of these alternative mechanisms. Ideally experiments of the type shown in Figure 2 could be used to validate the model, but at present the available data consists of histology images at two sparse time points (at the start and end of the experiments), whereas model validation and parameter estimation would require quantitative data at multiple time points. This is beyond the scope of the present study but could form the basis for future work in which the mathematical model here is used to identify the time points at which the data is collected.

**Figure 2:**
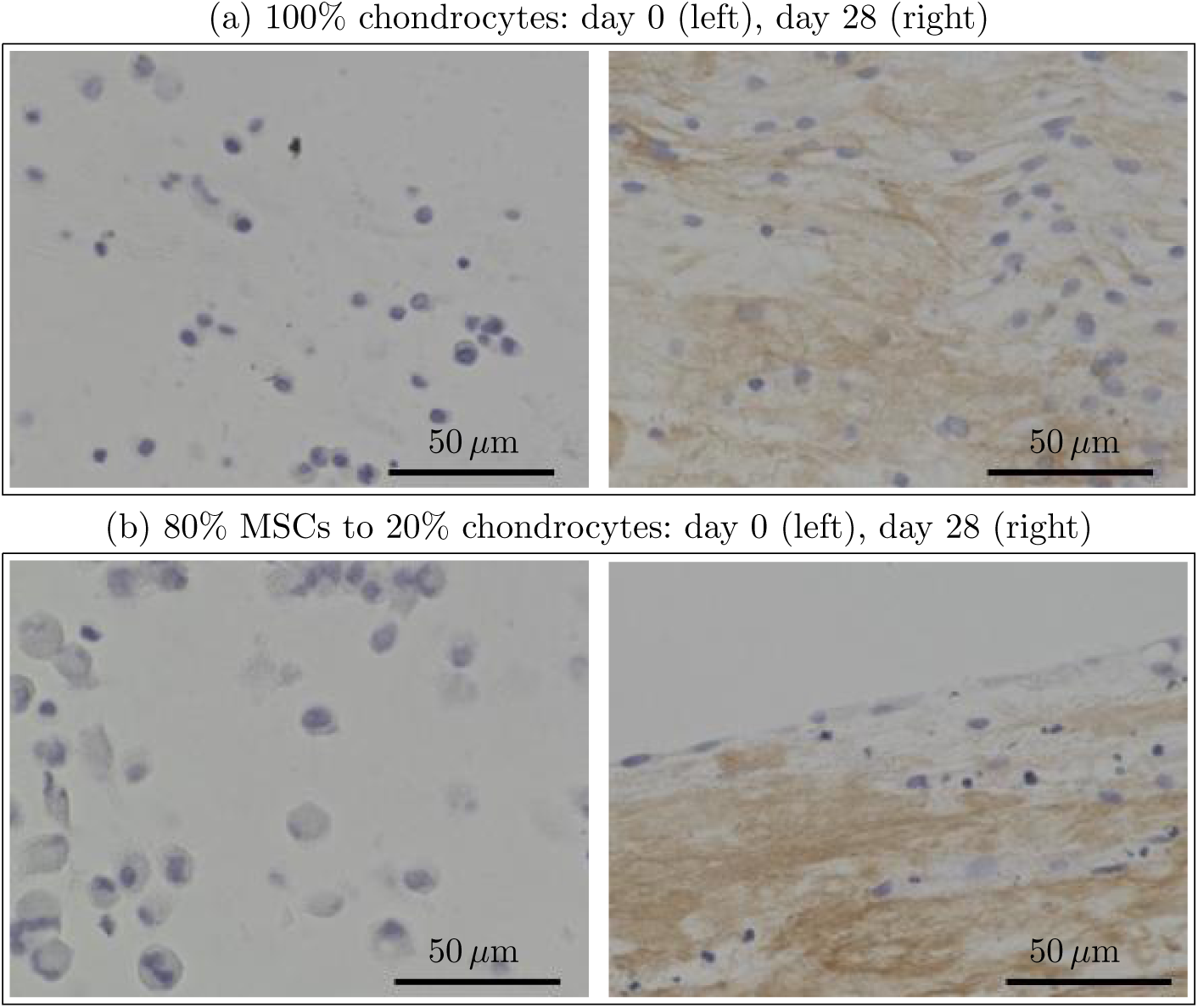
Microscope images from two *in vitro* experiments performed at University Hospital Wurzburg where a population of 100% chondrocytes and a co-culture of 80% MSCs and 20% chondrocytes were cultured for 28 days. In these images cells are stained in blue and type II collagen, which is produced by chondrocytes, is stained in brown. As shown in the images on the left in Figs. 2(a) and (b), no collagen is present in either culture at day 0. By contrast, the images on the right show in Figs. 2(a) and (b) show that at day 28 similar levels of type II collagen are present in both cultures. This indicates that the MSCs in the co-culture have differentiated into chondrocytes and are producing collagen type II. The scale bar in the lower right hand corner of each image is 50 *μ*m.

The purpose of this paper is to develop a mathematical model that describes the impact of TGF-*β* and chondrocytes on MSC proliferation and differentiation under various experimental conditions [8, 12, 13], and in doing so to identify the contributions from the various cellular and biochemical processes involved. A qualitative comparison is appropriate since the reported results in experiments are largely qualitative in nature, typically based on stainings for ECM components of the type shown in Fig. 2, and therefore not suitable when making quantitative comparisons. In formulating our model we adopt a highly simplistic view of the relevant biology, assuming well-mixed populations of cells and spatially uniform chemical concentrations, as well as simplifying aspects of the TGF-*β* life cycle, while retaining all the key mechanisms involved in chondrogenesis. Previous models have considered TGF-*β* decay and activation in the context of fibroblast differentiation [14], with a detailed focus on receptor kinetics, as well as modelled in detail the TGF-*β* signalling pathway [15]. We do not attempt to include this level of detail in our model, instead focussing on the interactions between extracellular TGF-*β* and the two cell types.

The remainder of this paper is organised as follows. In Section 2, we develop our mathematical model and present some motivating simulations. The chondrogenesis strategy of co-culturing MSCs and chondrocytes is investigated in Section 3, where we introduce a fast kinetics approximation and derive an analytic expression for the critical density of chondrocytes required to drive chondrogenesis. The effect of adding exogenous TGF-*β* on a population of cells is explored by considering early-time behaviour in Section 4, and we obtain the critical concentration of added TGF-*β* required to drive chondrogenesis with a combination of analytic and numerical techniques. Finally, a hybrid strategy where exogenous TGF-*β* is added to a co-culture is investigated in Section 5 with a combination of analytic and numerical techniques. We discuss these results and present our conclusions in Section 6.

### 2. Model and biological background

We develop a time-dependent mathematical model for TGF-*β* mediated proliferation and differentiation of MSCs to chondrocytes. We neglect cell motility and any spatial variation in the densities. We assume that chondrocytes do not proliferate or dedifferentiate, which is a reasonable assumption for tissue engineering applications, where the chondrocytes are typically derived from articular cartilage and cultured *in vitro*[16, 17, 18]. In this context it is also reasonable to neglect cell death of both MSCs and chondrocytes on the relatively short time scale of interest (approximately 1–2 weeks) [18].

We propose an ordinary differential equation model to investigate how the cytokine, TGF-*β*, can drive chondrogenesis in a well-mixed (spatially uniform) population of cells; the dependent variables of this model are summarised in Table 1 along with their units. We distinguish two cell types, MSCs and chondrocytes, and denote their densities by *m*(*t*) and *n*(*t*) (number of cells/volume), respectively. It is assumed that MSCs differentiate and proliferate at rates which depend on the amount of TGF-*β* bound to their receptors, denoted by *f* (*t*) (fraction of bound receptors/cell), with differentiation and proliferation occurring only if *f* (*t*) exceeds the threshold values *fd* and *fp*, respectively. Applying the principle of mass balance to the two cell types, the differential equations governing the evolution of *m*(*t*) and *n*(*t*) are

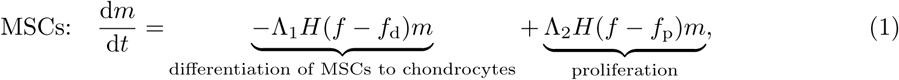

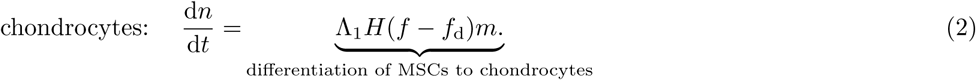

In Equations (1)–(2), *H*(*x*) is a Heaviside step function (*H(x)* = 1 if *x >* 0; *H(x)* = 0 otherwise), so that differentiation and proliferation are switched on when *f > f*d and *f > f*p, respectively. This represents a biologically appropriate description of the cell response to receptor bound TGF-*β*; following [19], for instance, the choice of Heaviside functions here has the advantage that it will permit further mathematical analysis. The constants Λ1 and Λ2 are the rates at which differentiation and proliferation, respectively, occur where these effects are active. Here we assume that *f*_*d*_ *< f*_*p*_, so that proliferation only occurs if the MSCs are also differentiating; this is broadly in line with the observed behaviour of MSCs that have been cultured *in vitro*, in a hydrogel, for a period of 1–2 weeks in the presence of the isoform TGF-*β*1 [5, 18, 20, 21, 22] in tissue engineering studies. We note that in other contexts and for other isoforms of TGF-*β* the effect may be different; for instance TGF-*β* in increases the proliferation of MSCs in bone formation [23], but has no effect on the proliferation of genetically engineered stem cells [24]. We further assume that Λ_1_ *>* Λ_2_, so that when both effects are active, proliferation does not outpace differentiation. This is in line with observations of the relative rates of these effects in experimental studies [20, 18]. We note that when Λ_1_ *<* Λ_2_ the model is physically unrealistic as the MSC population grows without bound.

In practice TGF-*β* can exist in up to eight different forms [9]. For simplicity, we distinguish just three forms of TGF-*β* and also track the quantity bound to the MSCs and chondrocytes. TGF-*β* is secreted as a large, latent molecular complex, which comprises the TGF-*β* ligand, a peptide which prevents interaction with cell receptors, and a protein which allows the complex to bind to the ECM. For our purposes latent TGF-*β* can exist in two states; an unbound form, which we denote by *c*(*t*) (mass/volume), and an ECM-bound form, which we denote by *b*(*t*) (mass/volume). In order for the ligand to be accessible to the cell receptors it must be activated, that is cleaved from the latent complex: we denote the concentration of active TGF-*β* by *a*(*t*) (mass/volume). A schematic diagram illustrating this life cycle is depicted in Figure 3.

**Figure 3:**
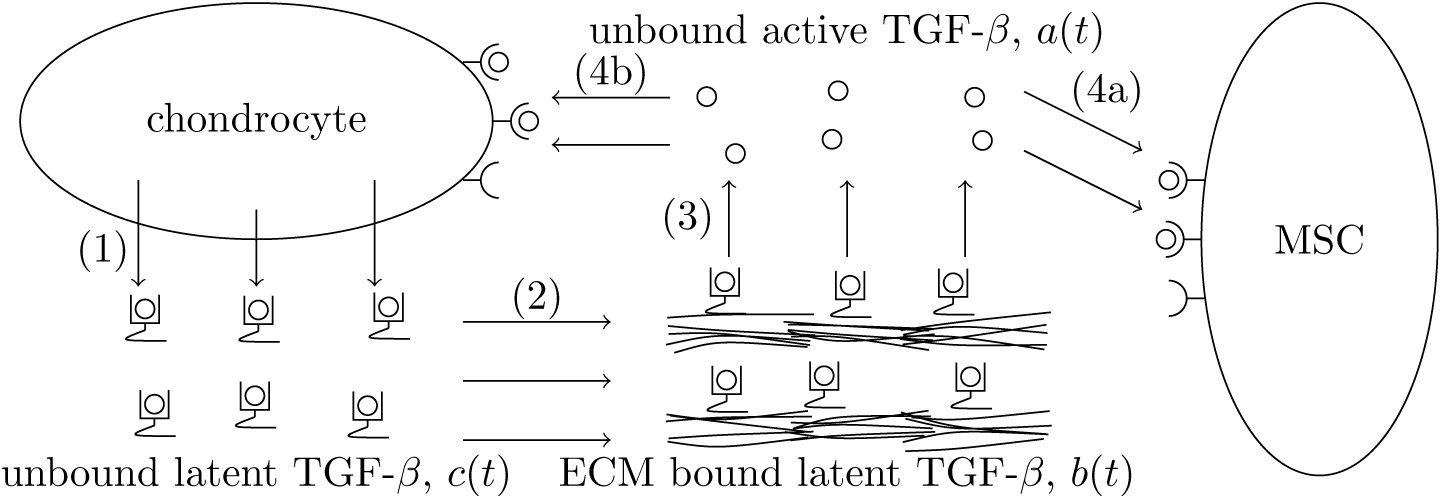
A schematic showing the simplified life cycle of TGF-*β* used in the model. The steps in this life cycle are: (1) secretion of latent TGF-*β* by chondrocytes; (2) binding of latent TGF-*β* to the extracellular matrix; (3) activation of bound TGF-*β* by interaction with an activating chemical species; (4a,b) binding of active TGF-*β* to the receptors on MSCs and chondrocytes.

Activation can arise from mechanical interaction with cell integrins [10] or through chemical interactions with proteases, thrombospondin-1, reactive oxygen species or extremes in pH [9]. Here, for simplicity, we assume that activation is caused by chemical interactions. Active TGF-*β* has a short half-life and so must bind rapidly to cell receptors or it will be degraded. As above, the fraction of bound TGF-*β* receptors per MSC for all cells in the population is denoted as *f* (*t*), and the fraction of bound receptors per chondrocyte for all cells in the population is denoted by *g*(*t*). Here, receptor-ligand dynamics are much simplified, as compared to other models [14, 15], since our interest is in cell differentiation rather than the details of these dynamics. We assume that when an MSC differentiates the bound TGF-*β* remains bound and, therefore, MSC differentiation will give rise to a loss of MSC-bound TGF-beta and an equal and opposite source of chondrocyte-bound TGF-beta. We further assume that each MSC or chondrocyte has a fixed number of receptors; so that if all receptors are occupied the total bound mass of TGF-*β* per MSC is denoted *F*_tot_, and the equivalent quantity for the chondrocytes is denoted *G*_tot_. The receptor bound TGF-*β* for both cell types is internalised at a constant rate, and we assume that when a ligand-receptor complex is internalised it is replaced by a free receptor.

As for the cell densities, we assume that the system is well-mixed so that the time evolution of the forms of TGF-*β* can be described by time-dependent ordinary differential equations (ODEs). The governing equations, along with a description of each term in underbraces, are presented in turn below:

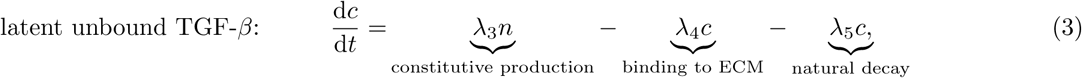

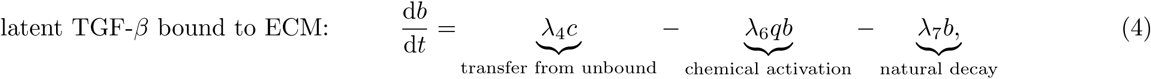

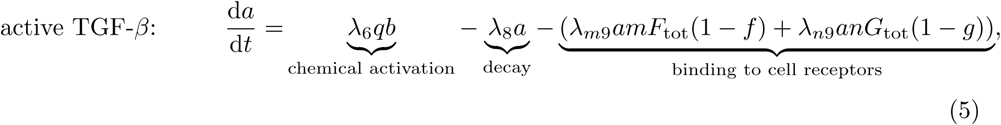

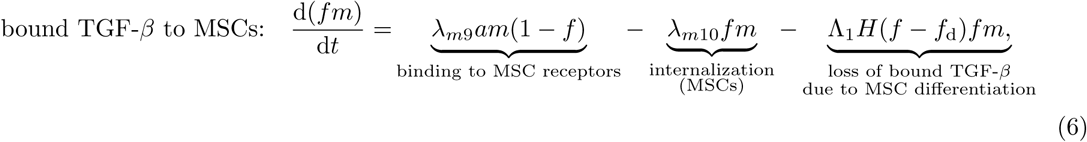

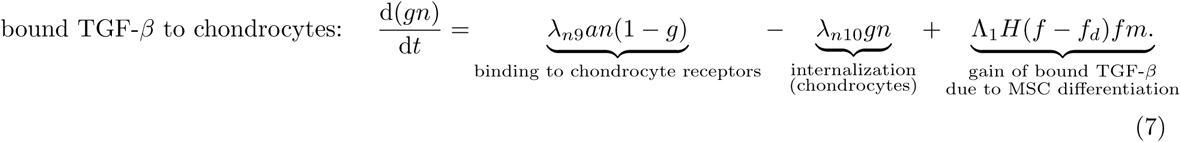

**Table 1:** Description of the model variables, along with their units. Here “Mio” is an abbreviation of “million”.

| Dependent variable | Description | Units |
| --- | --- | --- |
| $m$ | density of mesenchymal stem cells (MSCs) | Mio/mL |
| $n$ | density of chondrocytes | Mio/mL |
| $c$ | concentration of unbound, latent TGF-β | ng/mL |
| $b$ | concentration of bound, latent TGF-β | ng/mL |
| $a$ | concentration of unbound, active TGF-β | ng/mL |
| $f$ | fraction of bound TGF-β receptors per MSC | - |
| $g$ | fraction of bound TGF-β receptors per chondrocyte | - |

The dependent variables in Equations (1)–(7) are listed in Table 1 while descriptions of the model parameters, along with typical values, are given in Table 2. Most of these parameter estimates are taken directly from a combination of experimental studies and previous models of systems involving TGF-*β* [14, 15]. Others are estimated from experimental studies in combination with values from other studies. For example, the estimate of *Λ*_3_, is obtained from data given in [16] using parameter values from [25]. Similarly, in the absence of data with which to estimate the rate of differentiation, Λ_1_ the value in Table 2 was chosen to be consistent with the time scales over which *in vitro* studies of these phenomena are typically conducted. Estimates are not given for the proliferation rate Λ_2_ or the threshold values, *f*_*d*_ and *f*_*p*_; determining realistic values of these parameters is an aim of the present study. Note that the ‘binding to cell receptor’ terms in Equation (5) are multiplied by *F*_tot_ and *G*_tot_, whilst the corresponding terms in Equations (6) and (7) are not. This is because the units of *f m*, *gn* and *a* are different, and to ensure that all the terms in Equation (5) have the same units the ‘binding to cell receptors’ are multiplied by the maximum receptor bound TGF-*β* for each cell type.

**Table 2:** Summary of dimensional parameters that appear in Equations (1)–(7).

| Parameter | Description | Representative value |
| --- | --- | --- |
| $\Lambda_1$ | MSC to chondrocyte differentiation rate | $1.4 \times 10^{-3}/\text{min}$ |
| $\Lambda_2$ | MSC proliferation rate | - |
| $f_d$ | MSC to chondrocyte differentiation threshold | - |
| $f_p$ | MSC proliferation threshold | - |
| $\lambda_3$ | rate of constitutive TGF-β production by chondrocytes | $1.094 \times 10^{-1} \text{ ng}/(\text{min Mio})$ [16] |
| $\lambda_4$ | transfer rate between bound and unbound latent TGF-β | $10^{-1}/\text{min}$ [25] |
| $\lambda_5$ | decay rate of unbound latent TGF-β | $6 \times 10^{-3}/\text{min}$ [26] |
| $\lambda_6 q$ | activation rate due to chemical interactions | $6 \times 10^{-2}/\text{min}$ [16] |
| $\lambda_7$ | decay rate of bound latent TGF-β | $6 \times 10^{-3}/\text{min}$ [26] |
| $\lambda_8$ | decay rate of active TGF-β | $0.258/\text{min}$ [26] |
| $\lambda_{m9}$ | binding rate of active TGF-β to MSC receptors | $6 \times 10^{-2}/(\text{min ng/mL})$ [27] |
| $\lambda_{n9}$ | binding rate of active TGF-β to chondrocyte receptors | $6 \times 10^{-2}/(\text{min ng/mL})$ [27] |
| $\lambda_{m10}$ | internalisation rate of TGF-β bound MSC receptors | $0.25/\text{min}$ [15] |
| $\lambda_{n10}$ | internalisation rate of TGF-β bound chondrocyte receptors | $0.25/\text{min}$ [15] |
| $F_{\text{tot}}$ | maximum receptor bound TGF-β per MSC | $6.23 \times 10^{-1} \text{ ng}/\text{Mio}$ [28] |
| $G_{\text{tot}}$ | maximum receptor bound TGF-β per chondrocyte | $6.23 \times 10^{-1} \text{ ng}/\text{Mio}$ [28] |

The full model comprises Equations (1)–(7) along with appropriate initial conditions, which are slightly different for each of the chondrogenesis strategies. The existence and uniqueness of solutions to this model is an interesting open question, but one which we postpone for future work. The governing equations are non-dimensionalised by scaling the cell densities and concentration of TGF-*β* as follows

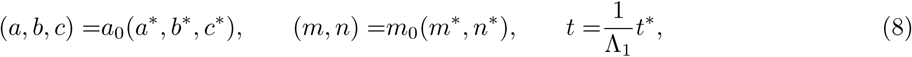

where stars denote dimensionless quantities. Here *a*_0_ and *m*_0_ are typical values for the concentration of TGF-*β* and cell densities; reasonable values for these scales, in the context of tissue engineering [3], are *a*_0_ = 1 ng/mL and *m*_0_ = 1 Mio/mL. The timescale of interest 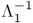 is that associated with cell differentiation; we shall see that the TGF-*β* dynamics operate on a much faster timescale than the timescale for differentiation. Under these scalings, Equations (1)–(7) become

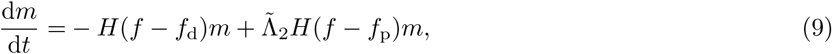

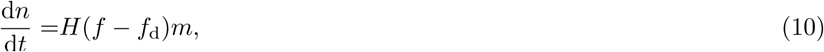

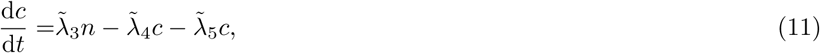

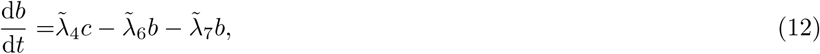

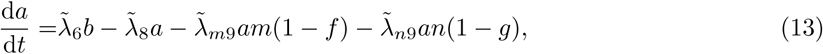

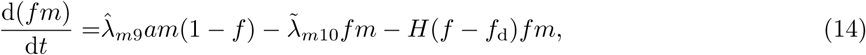

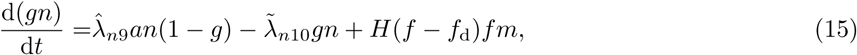

where we have dropped stars, and expressions for the dimensionless parameters are given in Table 3, along with representative values. Note that many of these parameters are much greater than 1, indicating that the dynamics of the transfer between the various forms of TGF-*β*, or their decay/internalisation, are much faster than MSC proliferation and/or differentiation. We exploit this separation of time scales between the chemical and cell processes by introducing the inverse of the decay rate of the active form as small parameter, 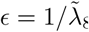, and scale the remaining large parameters as

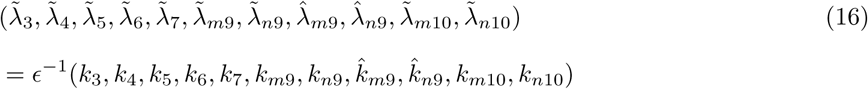

**Table 3:** Dimensionless modelling parameters appearing in the ODEs (9)–(15), together with values calculated from the dimensional parameter values in Table 2.

| Quantity | Definition | Value | Quantity | Definition | Value |
| --- | --- | --- | --- | --- | --- |
| $\tilde{\Lambda}_2$ | $= \Lambda_2/\Lambda_1$ | $\approx 1$ | $\tilde{\lambda}_{m9}$ | $= m_0 F_{\text{tot}} \lambda_{m9}/\Lambda_1$ | 6.23 |
| $\tilde{\lambda}_3$ | $= m_0 \lambda_3/(a_0 \Lambda_1)$ | 18.2 | $\tilde{\lambda}_{n9}$ | $= m_0 G_{\text{tot}} \lambda_{n9}/\Lambda_1$ | 6.23 |
| $\tilde{\lambda}_4$ | $= \lambda_4/\Lambda_1$ | 16.7 | $\hat{\lambda}_{m9}$ | $= a_0 \lambda_{m9}/\Lambda_1$ | 10 |
| $\tilde{\lambda}_5$ | $= \lambda_5/\Lambda_1$ | 1 | $\hat{\lambda}_{n9}$ | $= a_0 \lambda_{n9}/\Lambda_1$ | 10 |
| $\tilde{\lambda}_6$ | $= q \lambda_6/\Lambda_1$ | 10 | $\tilde{\lambda}_{m10}$ | $= \lambda_{m10}/\Lambda_1$ | 41.7 |
| $\tilde{\lambda}_7$ | $= \lambda_7/\Lambda_1$ | 1 | $\tilde{\lambda}_{n10}$ | $= \lambda_{n10}/\Lambda_1$ | 41.7 |
| $\tilde{\lambda}_8$ | $= \lambda_8/\Lambda_1$ | 43 | | | |

Substituting the rescaled parameters into (9)–(15) we obtain

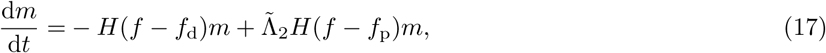

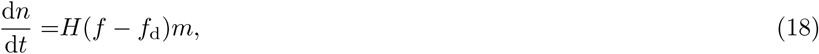

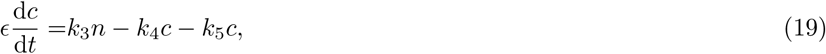

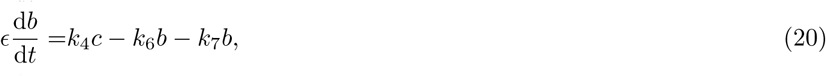

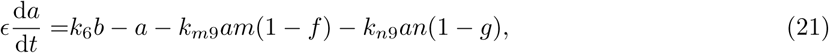

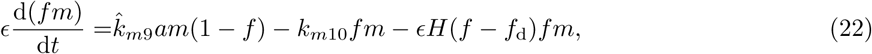

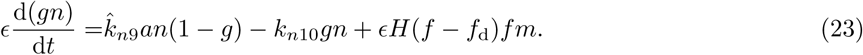

We note that the time derivatives in (19)–(23) are multiplied by the small parameter *ε*, reflecting that these dynamics occur on a fast time scale, while the cell processes represented by (17)–(18) act on a slower time scale.

### 2.1. Simulations of the full model

In the analysis that follows we shall see that the separation of time-scales between the chemical and cell processes leads to a rich solution structure. We begin by presenting numerical simulations of the full system (17)–(23) that typify common experimental chondrogenesis strategies; all simulations were performed in MATLAB with the Runge-Kutta (4,5) ordinary differential equation solver `ODE45`.

#### 2.1.1. Strategy 1: Co-culturing MSCs and chondrocytes to drive differentiation

One common differentiation strategy involves co-culturing MSCs and chondrocytes. Simulations for an initial MSC density of *m*(0) = 1 and two different initial chondrocyte densities are shown in Figure 4, with the initial concentrations of all forms of TGF-*β* zero (*c*(0) *= b*(0) *= a*(0) *= f* (0) *= g*(0) *=* 0). For the smaller initial density *n*(0) = 0.1 (represented by solid lines in Figure 4), TGF-*β* is constitutively produced by the chondrocytes and the concentrations of all forms of TGF-*β* rapidly attain a steady state at which TGF-*β* production is balanced by its natural decay and internalisation by the cells. As indicated in Figure 4(d) at steady state, and indeed for all times, the amount of TGF-*β* bound to the MSCs is below the critical value *f*_*d*_ at which differentiation occurs, and so no extra chondrocytes are produced meaning that this steady state persists for all time.

**Figure 4:**
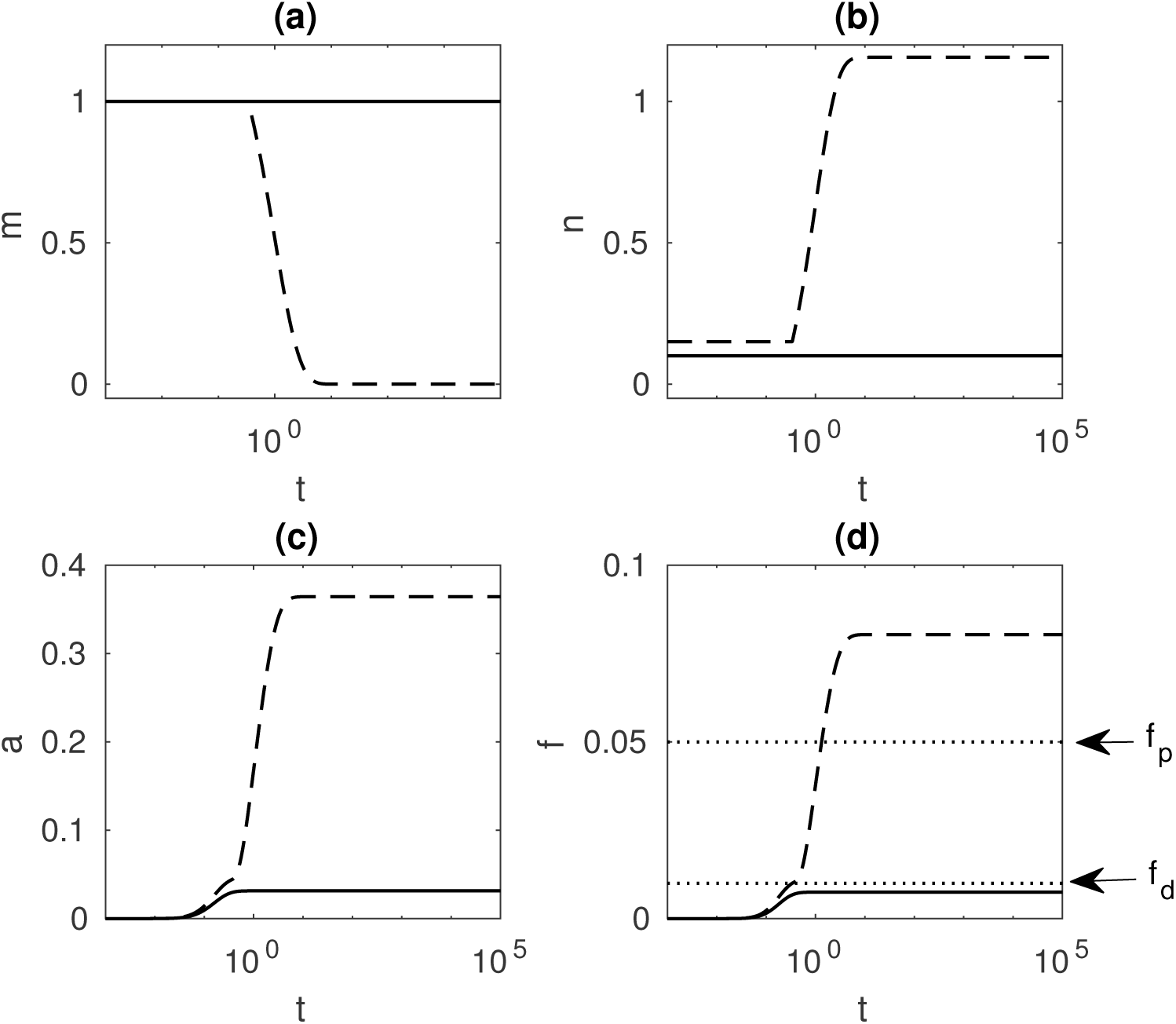
Numerical simulations of Equations (17)–(23) for selected components, using strategy 1 (chondrogenesis induced by co-culturing MSCs and chondrocytes). The *t*-axes in all these plots use a log scale to show both the short and long time behaviour clearly. In Figure 4(d) the dotted lines are for the critical values of *f* above which differentiation (*f*_*d*_ = 10^-2^) and proliferation (*f*_*p*_ = 5*×* 10^-2^) occur and the other parameters used in this simulation are given in Table 3. In Figure 4(a)–(d) the solid lines represent the case where *n*(0) = 0.1 and the dashed lines represent the case where *n*(0) = 0.15. For the smaller initial chondrocyte density the value of *f* never reaches *f*_*d*_ and so no MSC differentiation occurs. For the slightly larger initial chondrocyte density *f* exceeds *f*_*d*_ after a short time, so the whole initial population of MSCs is eventually differentiated, and *f > f*_*p*_ shortly thereafter so there is also proliferation of MSCs here.

For a slightly larger initial population of chondrocytes (*n*(0) = 0.15; dashed lines in Figure 4), the amount of TGF-*β* produced by the seeded chondrocytes is sufficient that the value of *f* = *f*_d_ is reached after a short time, triggering cell differentiation. At later times TGF-*β* levels continue to rise due to constitutive production by the newly produced cells, so that for the rest of the simulation *f > f*_*d*_ and *f > f*_*p*_ meaning that both differentiation and proliferation of the MSCs are triggered. The stem cells continues to differentiate until they are depleted and the system comprises only chondrocytes.

The final yield of chondrocytes, *n*_*∞*_, depends on the initial density of seeded chondrocytes *n*(0) as shown in Fig. 4 and in more detail in Fig. 5(a). If the initial density is less than some critical value, *n*_crit_, then no MSC differentiation occurs, and so the final yield of chondrocytes is simply the initial density (*n*(0) = *n*_*∞*_). If the initial density exceeds *n*_crit_ then eventually all the MSCs differentiate, so that *n*_*∞*_ *≥ m*(0) + *n*(0). As shown in Fig. 4(a), as *n*(0) is increased beyond *n*_*∞*_ there is a commensurate increase in the final yield *n*_*∞*_ due both to the extra initial seeded chondrocytes and a small amount of MSC proliferation. Practical interest lies in determining the value of *n*_crit_, since there is no advantage in seeding with more chondrocytes than is necessary to get the system into a state where all MSC will differentiate, as well as in understanding how *n*_crit_ relates to other parameters in the model. For example, as shown in Fig. 5(b) the value of *n*_crit_ increases as the threshold value for differentiation *f*_*d*_ is increased.

#### 2.1.2. Strategy 2: Exogenous TGF-*β* driving MSC differentiation

We now consider the case where exogenous TGF-*β* is added to a population of MSCs. Here, the initial cells densities are *m*(0) = 1 and *n*(0) = 0, so the initial population is purely MSCs, *a*(0) ≠ 0, representing the added TGF-*β*, and all other forms of TGF-*β* are initially zero. Simulations for two different initial values of *a*, the active form of TGF-*β*, are shown in Figure 6. For the smaller initial concentration (*a*(0) = 1; solid curves) the exogenous TGF-*β* decays rapidly and becomes negligible after a very short time. During this short time enough TGF-*β* binds to the MSCs such that for some early time *f > f*_d_ and a small number of chondrocytes are produced. However, not enough chondrocytes are produced to drive differentiation of the entire population of MSCs, and so a steady state persists where the cell population is mostly MSCs and a few chondrocytes with low concentrations of all forms of TGF-*β*.

**Figure 5:**
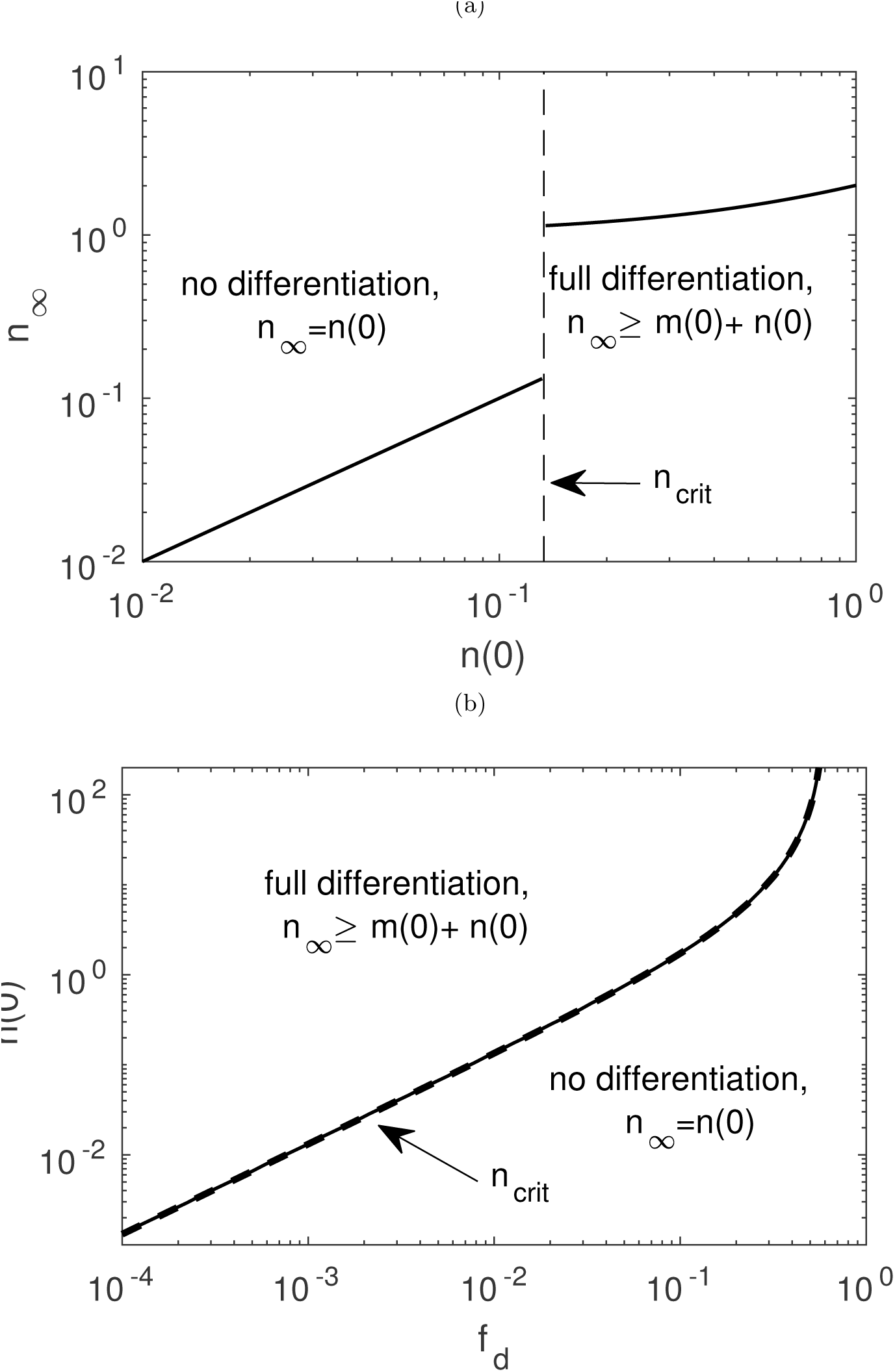
(a) The final yield of chondrocytes, *n_∞_*, for the co-culture strategy shown over a range of *n*(0), the initial density of added chondrocytes, as obtained from the numerical solutions of the full model (17)–(23). The vertical dashed line indicates *n*_crit_, the value of *n*(0) above which the constitutive production of TGF-*β* by the added chondrocytes is sufficient to drive complete differentiation. Here *f*_*d*_ = 10^-2^ and *f*_*p*_ = 5 *×* 10^-2^. (b) The critical value *n*(0) = *n*_crit_ shown against the modelling parameter *f*_*d*_, the threshold on TGF-*β* bound to MSC receptors above which differentiation occurs, with *f*_*p*_ = *f*_*d*_ + 4 *×* 10^-2^. The solid line is the value of *n*_crit_ obtained from the numerical solution and the dashed line is the analytic approximation to this value given in (31). The other parameter values used here are given in Table 3.

**Figure 6:**
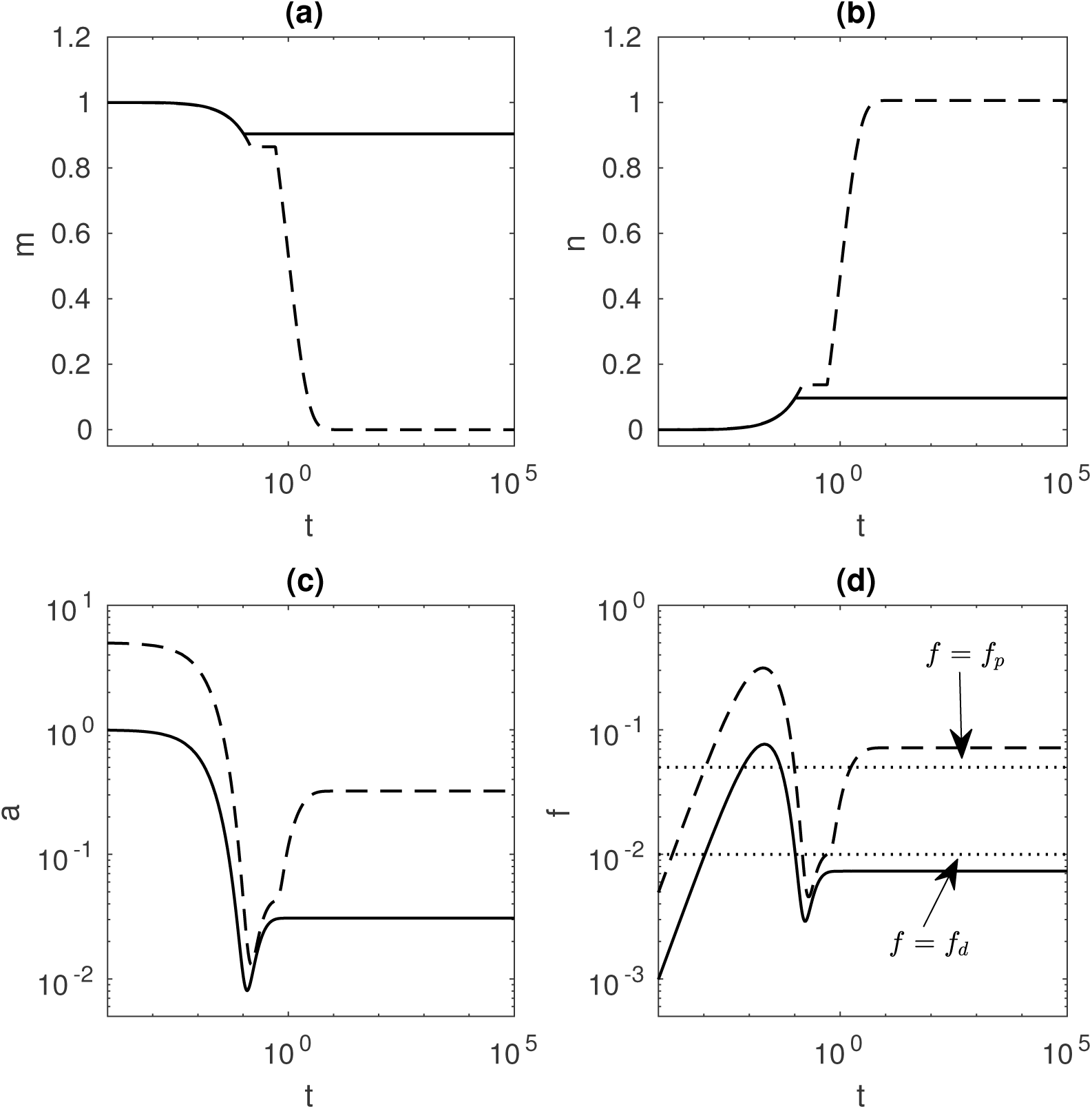
Numerical simulations of the full model (17)–(23) for selected solution components, for strategy 2 (chondrogenesis driven by exogenous TGF-*β*). The *t*-axes in all these plots use a log scale to show both the short and long time behaviour clearly. In Figure 6(d) the dotted lines indicate the critical values of *f* above which differentiation (*f*_*d*_ = 10^-2^) and proliferation (*fp* = 5 *×*10^-2^) occur; values of the other parameters used in this simulation are given in Table 3. For an initial active TGF-*β* concentration of *a*(0) = 1 (solid lines), there is incomplete chondrogenesis where only a fraction of the MSCs are differentiated. For an initial active TGF-*β* concentration of *a*(0) = 5 (dashed lines), sufficient chondrocytes are produced at early times to drive differentiation of the all of the MSCs plated in the dish at *t* = 0.

When the initial concentration of TGF-*β* is increased (from *a*(0) = 1 to *a*(0) = 5; dashed curves in Figure 6) more cells differentiate in the short time before the active TGF-*β* decays compared to the case where *a*(0) = 1. The larger chondrocyte population constitutively produces enough TGF-*β*(*f > f*_*d*_) to ensure all MSCs eventually differentiate. For completeness, we note that if a very low initial concentration of *a* is used, or *f*_d_ is large, then it is possible that no differentiation will occur.

The final chondrocyte yield *n*_*∞*_ is shown for a range of initial concentrations *a*(0) in Fig. 7(a), showing three possible outcomes. If the initial concentration is less than some critical value *a*_crit,1_, then the TGF-*β* degrades before any MSC differentiation takes place, giving a long time cell population consisting of the initial MSCs and no chondrocytes, and all forms of TGF-*β* zero. For moderate size concentrations, where *a*_crit,1_ *< a*(0) *< a*_crit,2_, a few cells will be differentiated at early times but these are insufficient to drive the entire population to differentiate; the long time cell population thus consists of mostly MSCs and a few chondrocytes, with low levels of all forms of TGF-*β*. For larger initial concentration, with *a*(0) *> a*_crit,2_, sufficient chondrocytes are produced at early times to ensure that eventually all MSCs differentiate. This second critical value is of great practical interest, since full differentiation of a MSC population is ultimately the desired outcome. The dependence of *a*_crit,1_ and *a*_crit,2_ on the differentiation threshold parameter *f*_*d*_ is shown in Fig. 7(b). Where *f*_*d*_ is small there are no initial concentrations for which partial differentiation is possible, and so if any differentiation occurs the system will be driven to full differentiation; where *f*_*d*_ is large there are no initial concentrations where full differentiation is possible. Experimental studies display both partial and full differentiation depending on the initial concentration (as well as situations where no chondrocytes are produced), which indicates that realistic values for *f*_*d*_ are in the range where all three of these outcomes can occur (for instance at *f*_*d*_ = 10^-2^ in Fig. 7(b)).

**Figure 7:**
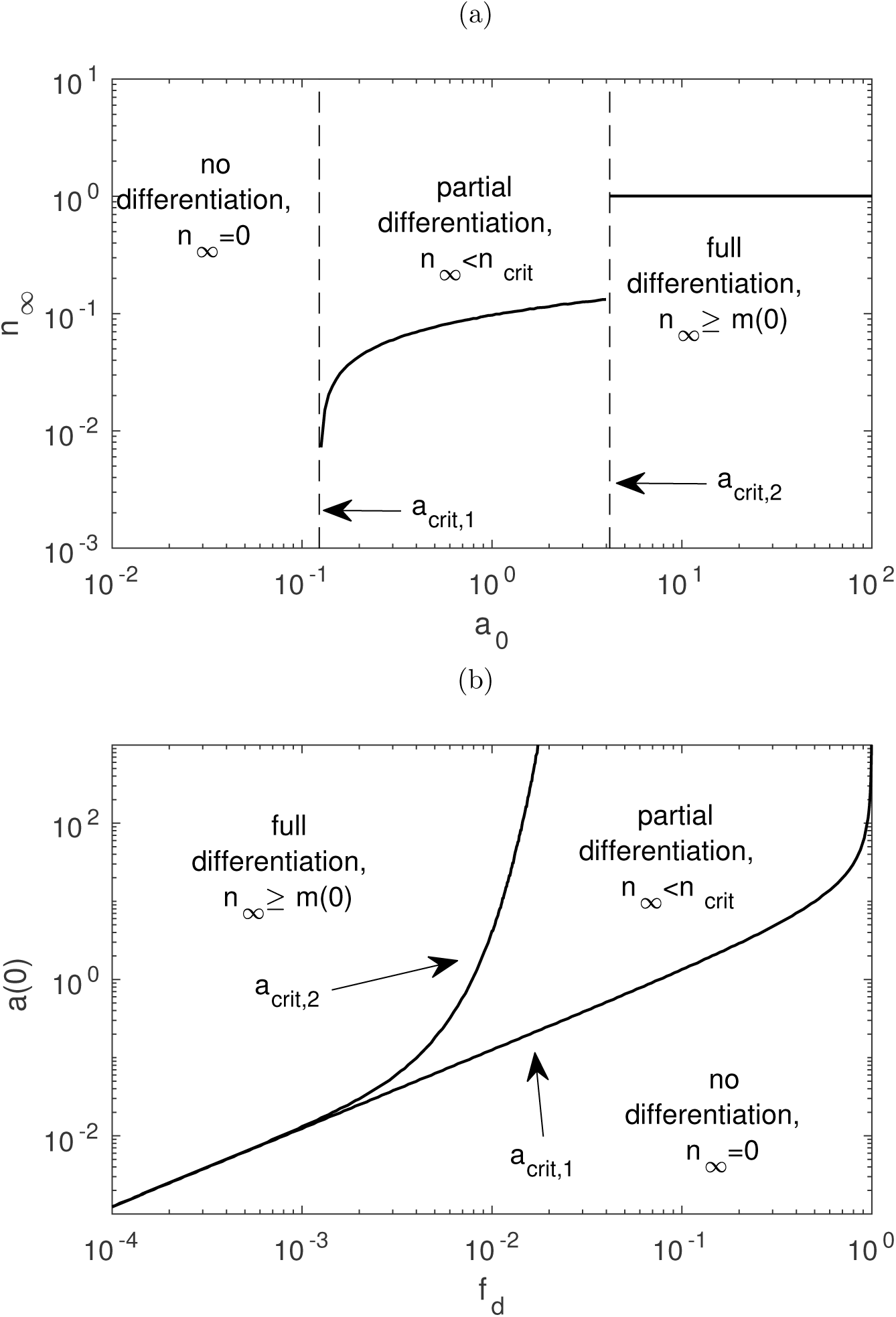
(a) The final yield of chondrocytes, *n_∞_*, for the strategy 2, over a range of *a*(0), the initial concentration of exogenous TGF-*β*, as obtained from the numerical solution of the full model (17)–(23). The left vertical dashed line indicates *a*_crit,1_, the value below which no chondrocytes will be produced, and the right vertical dashed line indicates *a*_crit,2_, the value above which complete differentiation occurs; between the these two critical values there is partial differentiation, and in this region the final yield *n*_*∞*_ increases with the initial concentration *a*(0). Here, *f*_*d*_ = 10^-2^, *f*_*p*_ = 5 *×* 10^-2^. (b) The critical values *a*_crit,1_ and *a*_crit,2_ shown against the modelling parameter *f*_*d*_, the threshold on TGF-*β* bound to MSC receptors above which differentiation occurs. Both critical curves are from the numerical solution to (17)–(23) and analytic approximations to these curves are obtained in Section 4. Here, *f*_*p*_ = *f*_*d*_ + 4 *×* 10^-2^. For both Fig. (a) and (b) all other parameter values are given in Table 3; thus the value of *n*_crit_ is the same as in Fig. 5.

## 3. Strategy 1: Chondrogenesis in co-cultures of MSCs and chondrocytes

The simulations presented in Figures 4 and 5 reveal that a population of MSCs can fully differentiate when they are co-cultured with a small number of chondrocytes if the initial proportion of chondrocytes exceeds some critical value, *n*_crit_ say. That is, for a given initial chondrocyte density *n*(0) two long term outcomes are possible:

- No differentiation: *n*(0) *< n*_crit_ *⇒ n*_*∞*_ = *n*(0);
- Full differentiation: *n*(0) *> n*_crit_ *⇒ n*_*∞*_ *≥ m*(0) + *n*(0).

In experimental studies the percentage of chondrocytes in co-cultures typically ranges from 25–66% [12], although ideally experimentalists aim to use fewer chondrocytes, since they are difficult to harvest [13]. Note that the chondrogenesis experiment in Fig. 2 used only 20% chondrocytes and the numerical simulations in Section 2.1.1 predicted that even fewer chondrocytes were required to drive differentiation, in turn this suggests that (assuming our parameter values are reasonable) fewer chondrocytes may need to be harvested to successfully employ this strategy.

We now exploit the different time scales of the cell process and TGF-*β* dynamics to make a quasi-steady (or ‘fast kinetics’) approximation to (19)–(23). This will allow us to estimate *n*_crit_ in terms of the model parameters.

### 3.1. Fast-kinetics approximation of the TGF-*β* dynamics

Since the TGF-*β* dynamics occur on a faster timescale than the cell processes it is appropriate to make a quasi-steady approximation to Equations (19)–(23) by taking the limit as ϵ *→* 0. Equations (19)–(23) then reduce to the following algebraic expressions relating the various components (where a subscript ‘FK’ denotes the quasi-steady, or ‘fast-kinetics’, approximation of a particular variable),

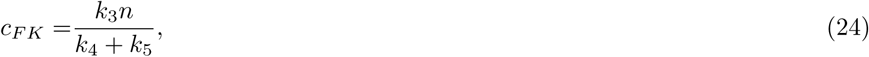

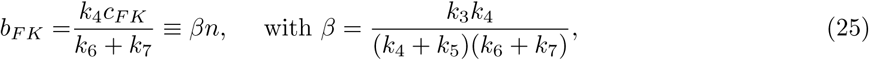

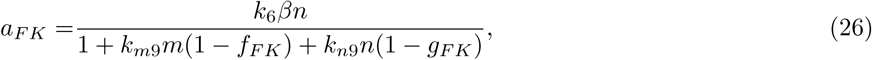

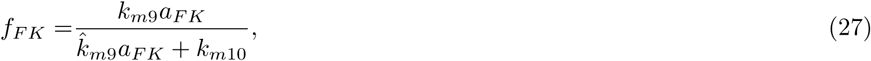

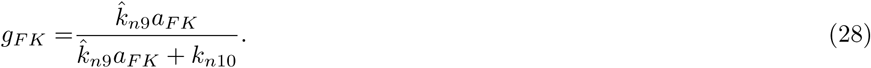

Equations (24)–(28) implicitly define the time-evolution of the variables *c*, *b*, *a*, *f* and *g*, via their dependence on *m* and *n* whose evolution is defined by Equations (17) and (18). From (24) and (25) we note that *c*_*F*__*K*_(*t*) and *b*_*F*__*K*_(*t*) are proportional to the chondrocyte density, *n*(*t*). The remaining variables *a*_*F*__*K*_(*t*), *f*_*F*__*K*_(*t*) and *g*_*F* *K*_(*t*) are interdependent; while no simple expression in terms of the cell densities exists, some further analysis is possible. Substituting expressions (25), (27) and (28) into (26) we obtain the following cubic equation for *a*_*F*__*K*_(*t*) = *a*_*F*__*K*_(*m*(*t*)*, n*(*t*)):

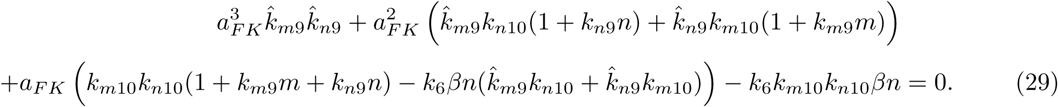

We note that for *n >* 0 this expression has one real positive root, since its discriminant is always positive, and two stationary points at which *a*_*F*__*K*_ *<* 0.

Following some straightforward manipulation of the fast kinetics approximation (24)–(28) we can determine *n*_crit_, the critical value for the initial concentration of chondrocytes above which differentiation of the whole population occurs. This is the value of *n* which gives *f*_*F*__*K*_ = *f*_d_, the threshold value of *f* above which differentiation occurs. This value of *f*_*F*__*K*_ is associated with a threshold value of *a*_*F*__*K*_ = *a*_*d*_, and from rearrangement of (27) we find

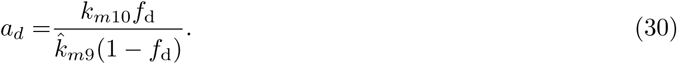

Substituting this value into equation (28) we define 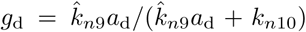. By rearrangement ofequation (26) we obtain *n*_crit_ in terms of *m*(0) and the system parameters, namely

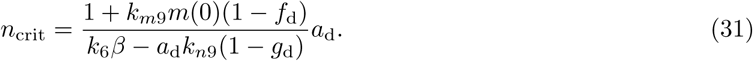

The value of *n*_crit_ in (31) is in excellent agreement with the value obtained from the numerical solution of the full governing equations (dashed curve in Fig. 5(b)). We note that (31) depends on all the system parameters given in Table 3, except for the proliferation rate Λ_2_. This analytic expression reveals the dependence of the critical value on the model parameters. For example, we note that an increase in *k*_6_, the rate at which the latent form of TGF-*β* is activated, leads to a commensurate decrease in *n*_crit_, reflecting that the constitutively produced TGF-*β* is being transferred more efficiently.

## 4. Strategy 2: Chondrogenesis induced by exogenous TGF-*β*

We now determine the conditions under which a population of MSCs will eventually be fully differentiated by adding exogenous TGF-*β*. Although any added TGF-*β* quickly decays, as seen in the numerical simulations in Figure 6, it may still have a significant effect on the system in this short time period. The route to chondrogenesis is that in this short time a few MSCs are differentiated, and the resulting chondrocytes drive the remaining MSCs ultimately to differentiate in the manner described in Section 3. The simulations in Section 2.1.2 revealed three possible outcomes arising from the addition of an initial concentration *a*(0) of active TGF-*β* to a population of pure MSCs:

- No differentiation: *a*(0) *< a*_crit,1_ *⇒ n*_*∞*_ = 0;
- Partial differentiation: *a*_crit,1_ *< a*(0) *< a*_crit,2_ *⇒* 0 *< n*_*∞*_ *< n*_crit_;
- Full differentiation: *a*(0) *> a*_crit,2_ *⇒ n*_*∞*_ *≥ m*(0).

We examine the behaviour of (17)–(23) at early times using perturbation series techniques and, in so doing, determine how many chondrocytes are produced in response to the exogenous TGF-*β*, which in turn allows us to approximate *a*_crit,1_ and *a*_crit,2_ for different values of the initial concentration *a*(0).

### 4.1. Early times

The short half-life of active TGF-*β* means that any MSC differentiation associated with exogenously added TGF-*β* occurs a short time after it has been introduced to the culture medium. To this end we now analyse Equations (17)–(23) at early times in response to the addition of a known amount of active TGF-*β* at *t* = 0. Our goal is to estimate how many chondrocytes (if any) are produced during this short time.

To determine whether differentiation and/or proliferation will occur we need to solve for *f*. Thus for this analysis it is convenient to use Equations (17) and (18) to rewrite Equations (22) and (23) as differential equations for the per cell quantities *f* and *g* rather than the quantity per volume *fm* and *gn*. These equations become

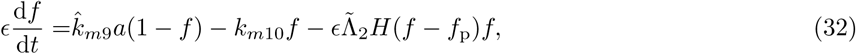

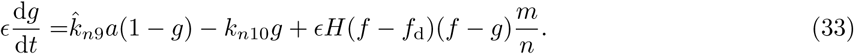

When active exogenous TGF-*β* is added to population of MSCs the initial conditions are

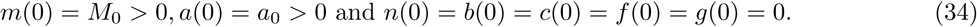

In general, these initial conditions will be incompatible with the fast-kinetics approximation (24)–(28), and so there is boundary layer at *t* = 0. We scale into this layer by setting *t* = ϵ*τ* and rescale *a* with a parameter *δ* that indicates the size of its initial magnitude, so that *a* = *δA* with *A ∼ 𝒪*(1) and *a*_0_ *∼ 𝒪*(*δ*); in Section 4.1.1 we take *δ* = ϵ to consider small initial concentrations of *a*, and then in Section 4.1.2 take *δ* = 1 to examine moderate initial concentrations of *a*. Substituting these scalings into (17)–(21), (32) and (33) gives

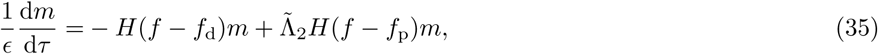

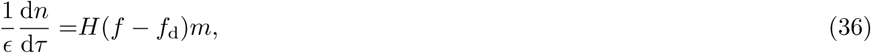

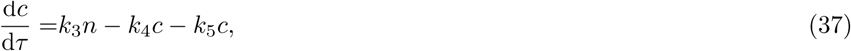

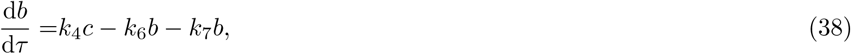

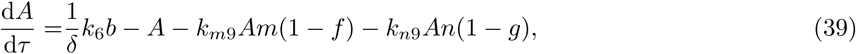

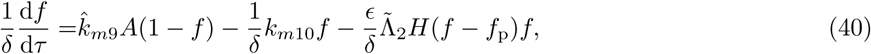

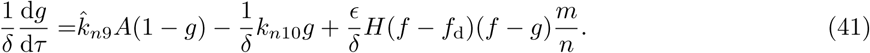

Perturbation series expansions in power of ϵ are introduced for each variable; these are of the form

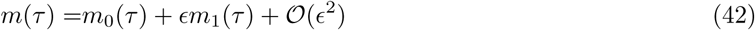

with similar expansions for *n*, *c*, *b*, *A*, *f* and *g*. We now examine equations (35)–(38), which do not involve the amplitude parameter *δ*, and will examine (39)– (41) (which do involve *δ*) separately in Sections 4.1.1 and 4.1.2 as the forms these equations take is different for small and moderate initial amplitudes of *a*. Substituting the approximate solutions (expansions) into (35)–(36), the leading order equations for the cell densities are

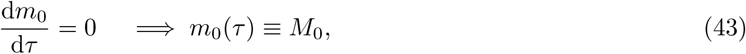

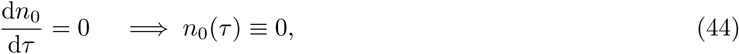

that is, at leading order, the cell densities do not change from their initial values. Continuing to 𝒪(ϵ) gives

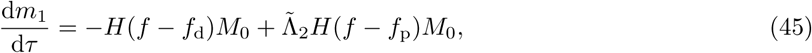

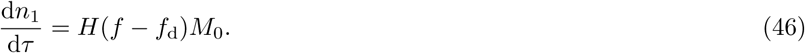

Our primary interest is in whether the exogenous TGF-*β* induces differentiation of MSCs at early times and, if so, to establish how many chondrocytes are produced; this is determined by integration of (46) as

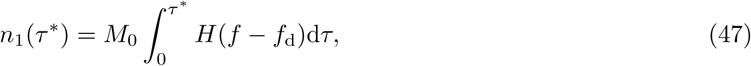

where *τ* ^*\**^ is an 𝒪(1) parameter for the width of the boundary layer (an appropriate choice for this is *τ*^*\**^ = 2 log(1*/*ϵ), namely the time at which an exponentially decaying initial concentration of *a* will have decayed by two order of magnitudes in ϵ). Similarly Equations (37) and (38) supply the following differential equations at leading order:

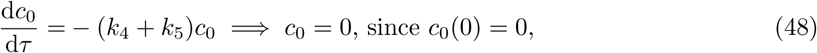

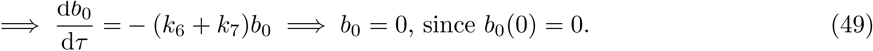

These leading order solutions are consistent with there being no chondrocytes at leading order, and therefore no constitutive production of latent TGF-*β*. The corresponding equations at 𝒪(ϵ) are

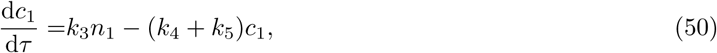

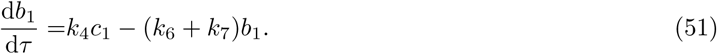

The leading-order (and first-order) forms of Equations (39)–(41) differ slightly depending on the choice of *δ*, the initial magnitude of *a*, and in the following analysis we consider the cases where the system is excited by different initial concentrations at *δ* = ϵ and *δ* = 1, which each reveal different aspects of the behaviour of the system.

#### 4.1.1. Small initial concentration: *δ* = *ϵ*, *a*(0) = ϵ*A*(0) *∼ 𝒪*(ϵ)

Here we assume the initial concentration is small, so that *a*(0) = ϵ*A*(0) = ϵ*α*_0_, where *α*_0_ *∼ O*(1). We seek to estimate *a*_crit,1_, the critical initial value of concentration *a*(0) below which no MSC differentiation occurs. To this end we assume that for these small initial concentrations no differentiation or proliferation occurs, that is *f < f*_d_ and *f < f*_p_, which implies that *n*_1_ = 0 and *m*_1_ = 0 from (45) and (46). By a similar argument to that applied to solve Equations (48) and (49), it follows from (50) and (51) that *c*_1_ = *b*_1_ = 0. The leading order equations for *f* and *A* are then

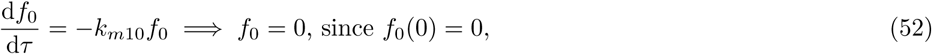

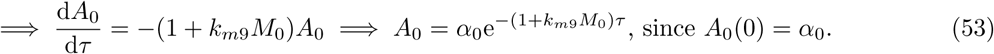

Continuing to 𝒪(ϵ) in *f* we have

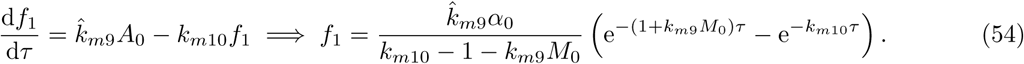

These solutions are in excellent agreement with the numerical solution of the full system. See, for example, Figure 8(a)–(b) where for *a*_1_ the relative error between the numerical solution and the approximate solution (53) is at most 1.76%, and for *f*_1_ the relative error between the numerical and approximate solution (54) is at most 3.64%.

It is straightforward to show that *f*_1_ attains its maximum at some time 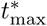. We seek the value of theinitial concentration *α*(0) = ϵ*α*_0_ at which this maximum corresponds to the critical value for differentiation (ie. *Ef*_1_ = *f*_d_). Rearranging (54), we deduce that no differentiation will occur if

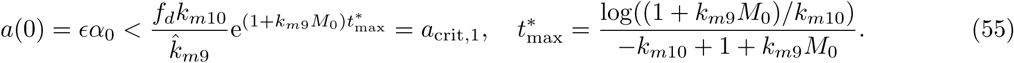

This inequality provides an asymptotic estimate for the first of the critical values in Figure 7(a), that is the value that separates regimes in which there is no cell differentiation from regimes in which there is partial differentiation. In Figure 8(c) we show that this approximation is in excellent agreement (less than 6.22% relative error where *a*(0) *<* 1) with the numerical results generated from the full model when *f*_*d*_ is small, and (as might be expected) underestimates the critical value in the limit as *f*_*d*_ *→* 1. Equation (55) reveals that this critical value depends only on *f*_*d*_ and the system parameters involved in the binding and internalisation of TGF-*β* to MSCs. This reveals, for instance, that *a*_crit,1_ increases with the internalisation rate *k*_*m*10_, indicating that more exogenous TGF-*β* must be added to achieve any cell differentiation if TGF-*β* is taken up by the cells at a higher rate.

To summarise, this analysis confirms the numerical result from Section 2.1.2 that the addition of a small concentration of TGF-*β* results in either no cells being differentiated or partial differentiation of the cell population. The key prediction is the analytic approximation of the cut-off TGF-*β* concentration given in Equation (55), below which no differentiation occurs, and the dependence of this value on various model parameters.

#### 4.1.2. Moderate initial concentration: *δ* = 1, *a*(0) = *α*_0_ *∼ 𝒪*(1)

When a moderate concentration of exogenous TGF-*β* is added to a culture medium there is typically some differentiation of cells at early times, as indicated in the simulations in Section 2.1.2. The analysis of the previous section provided an excellent approximation of *a*_crit,1_, and we now aim to estimate *a*_crit,2_, the critical concentration above which sufficient cells are produced to drive differentiation of the whole population. The simulations presented in Section 2.1.2 suggest that *a*_crit,2_ is approximately an order of magnitude larger than *a*_crit,1_. Therefore, in what follows we will assume that *a*_crit,2_ = 𝒪(1).

We assume that the *a*(0) = 𝒪(1) and *δ* = 1. Under these assumptions, it follows that at leading order Equations (39) and (40) supply

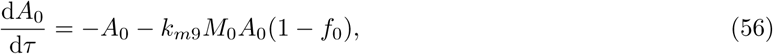

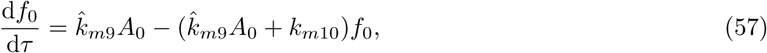

**Figure 8:**
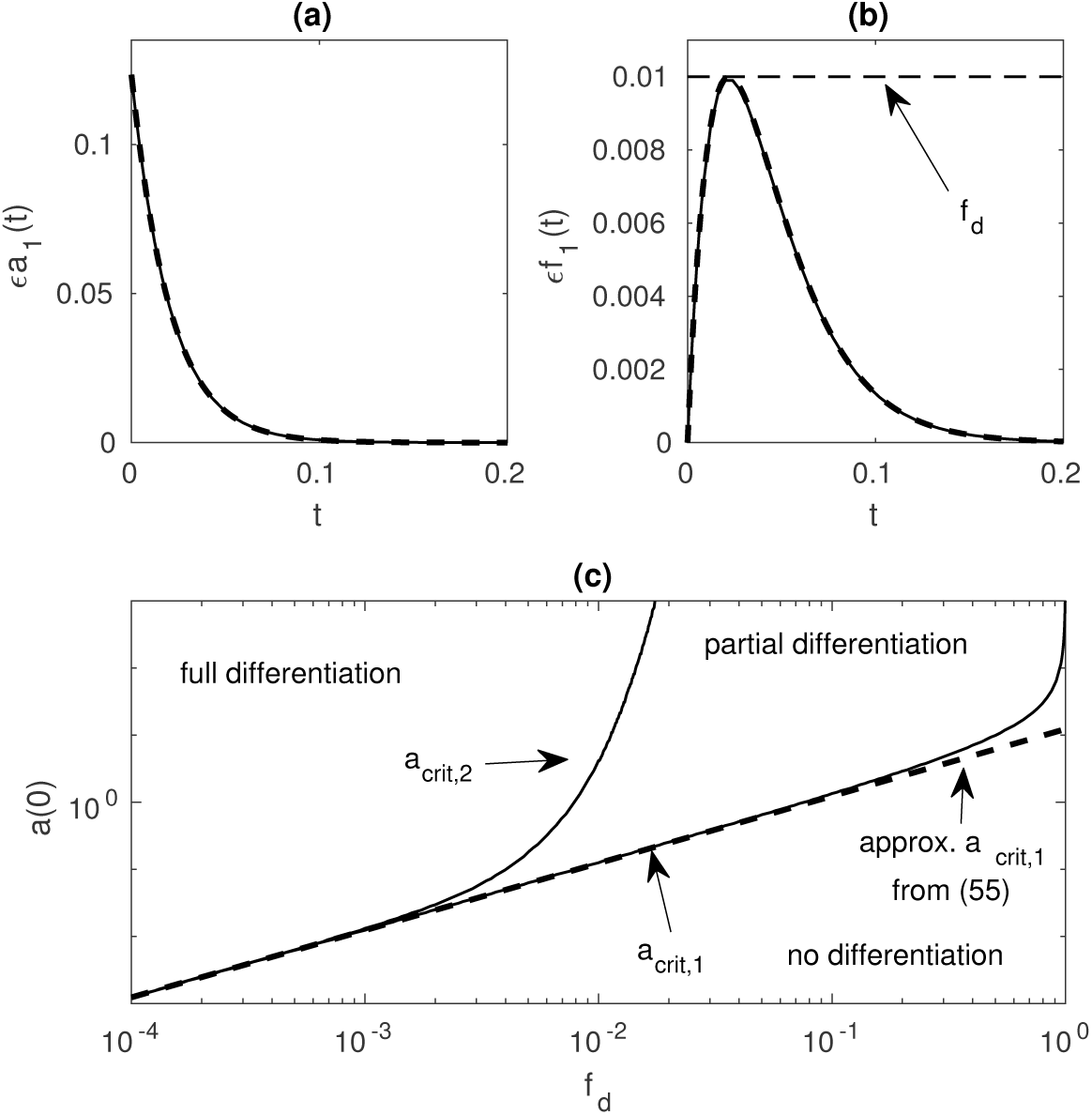
Early-time solutions for a small initial concentration of *a*. The solution components *a* = *εa*_1_(*t*) and *f* = *εf*_1_(*t*) are shown in Figures 8(a) and (b), respectively. The horizontal dashed line in Figures 8(b) is the threshold value *f*_*d*_ = 10^-2^. This solution for *f* reaches, but does not cross this threshold, indicating that the value of *a*(0) = *a*_crit_. The approximation to this critical value from (55) is shown as a dashed line in Figure 8(c) against the numerical critical values (solid lines) and is nearly indistinguishable for small *a*(0) and *f*_*d*_ *≪* 1. Here, *f*_*p*_ = *f*_*d*_ + 4 *×* 10^-2^ and all other parameter values are given in Table 3; thus the value of *n*_crit_ is the same as in Fig. 5.

**Figure 9:**
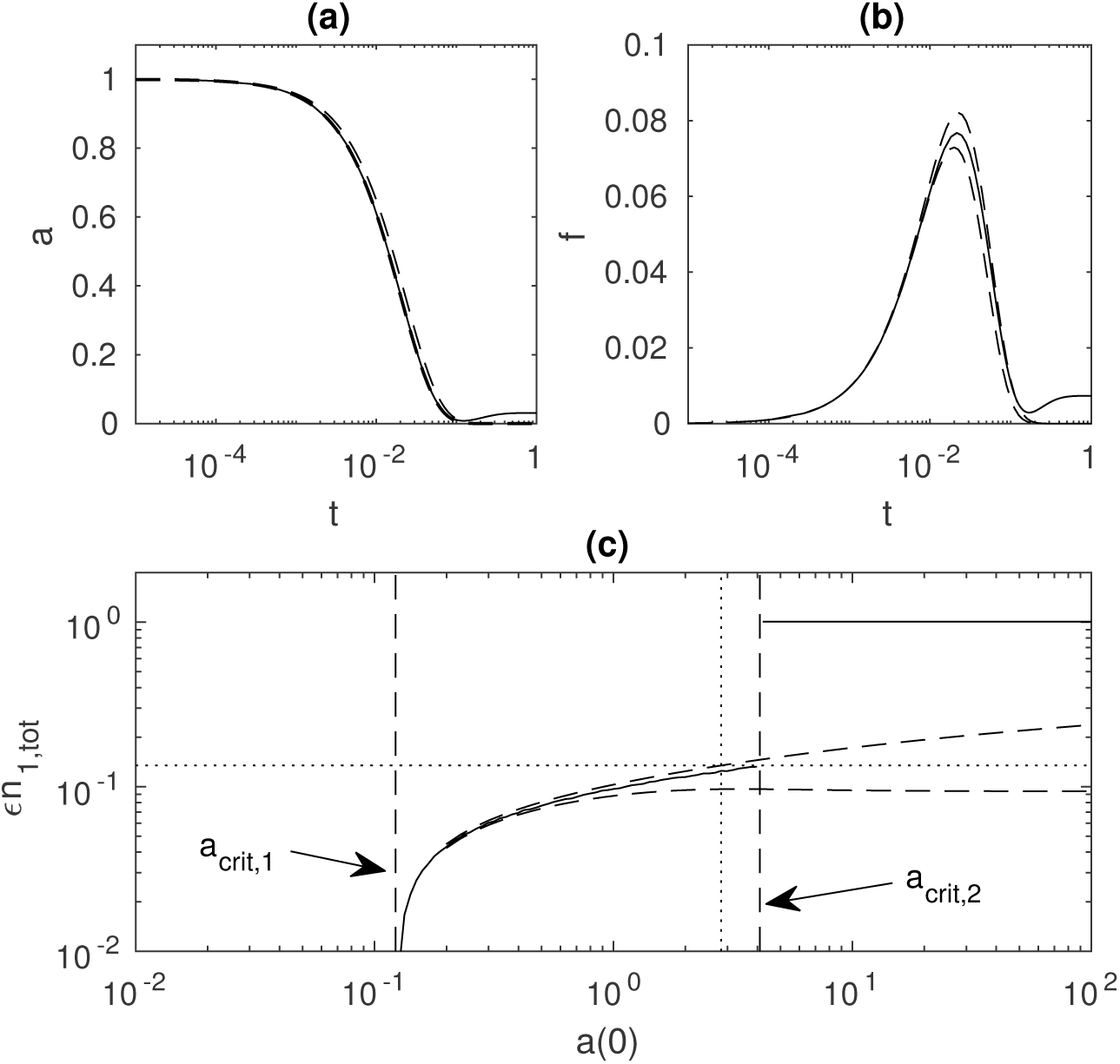
Early time solutions for a moderate initial concentration of *a*. (a) The numerical solution for *a*(*t*) (solid line) is bounded by the upper and lower approximations from (58) and (59), both shown as dashed lines (note here that the dashed line for the the lower approximation is partially obscured by the solid line). (b) The numerical solutions for *f* (*t*) (solid line) is bounded by the upper and lower approximations from (61) and (63), both shown as dashed lines. (c) The chondrocyte yield *n∞* (solid line) shown against initial concentration *a*(0), along with the predicted yield after early time from the upper and lower approximations (dashed curves). The vertical dashed lines are numerical values of *a*_crit,1_ and *a*_crit,2_. The horizontal dotted line is *n*_crit_ from (31), and the vertical dotted line is the approximate value of *a*_crit,2_ calculated from the upper bound. Note that in this case no approximation of *a*_crit,2_ from the lower bound is possible since the lower bound never reaches the critical value *n*_crit_. Here, *f_p_* = 5 *×* 10^-2^ and all other parameter values are given in Table 3.

Since both *b* and *n* are zero to leading order, and this system has no closed form solutions. It is possible, however, to bound the solution to *A*_0_ by using the fact that 0 *≤ f*_0_ *≤ F*_0_ in (56), where 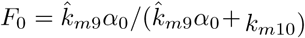 from the fast-kinetics expression (27); that is, as the initial concentration of *a* is decaying, we assume that *f* does not exceed the quasi-steady value it would take when the a concentration of *a* = *α*_0_ is present in the system. The upper and lower bounds on *A*_0_ are obtained by setting *f*_0_ = *F*_0_ and *f*_0_ = 0, respectively, in (56) which gives

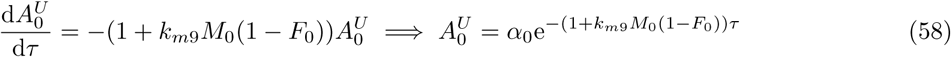

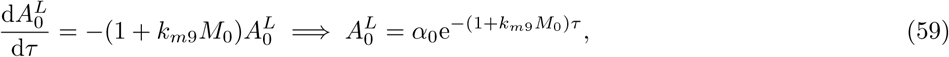

and the lower bound in particular is an excellent approximation to the numerical solution of *a*; see, for instance, Fig. 9(a) where the relative error between the approximation (59) and the numerical solution is less than 2% for *t <* 0.05. These bounds are then used in (57) to determine upper and lower bounds on *f*_0_. To find the upper bound on *f*_0_ we set 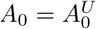 in the first term on the right hand side (so that the effect of this term is maximised) of (57) and *A*_0_ = 0 in the second term (so that the effect of this term is minimised) to give

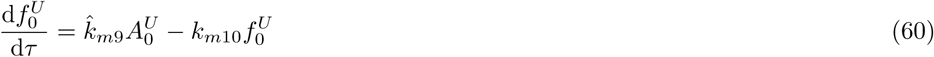

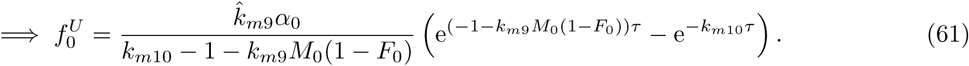

To obtain a lower bound we set the 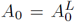 in the first term and *A*_0_ = *α*_0_ in the second term, thereby minimising the growth and maximising the decay of *f*_0_, so that (57) becomes

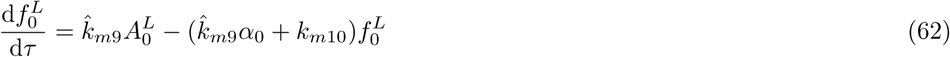

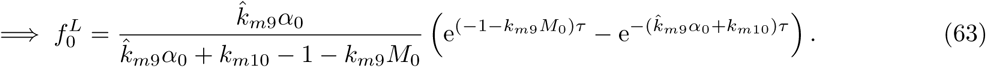

These bounds for *f*_0_ are compared with the numerical solution in Fig. 9(b) with the upper bound again being a particularly close approximation to the numerics with a relative error of at most 8.61% for *t <* 0.1.

Having bounded *f*_0_ we can estimate the number of chondrocytes produced in response to a given initial concentration of *a*. To determine an upper estimate of early time chondrocyte yield we determine the length of time for which 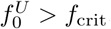, that is the time for which the step function in (47) is switched on. This time is estimated by finding the roots of 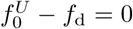, and noting that two such roots exist provided *a*(0) *> a*_crit,1_ from (55) (that is, when some MSCs have been differentiated). Having obtained these two values, 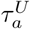 and 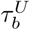, it follows from (47) that the number of differentiated cells is

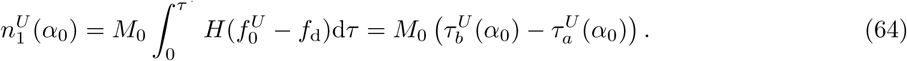

**Figure 10:**
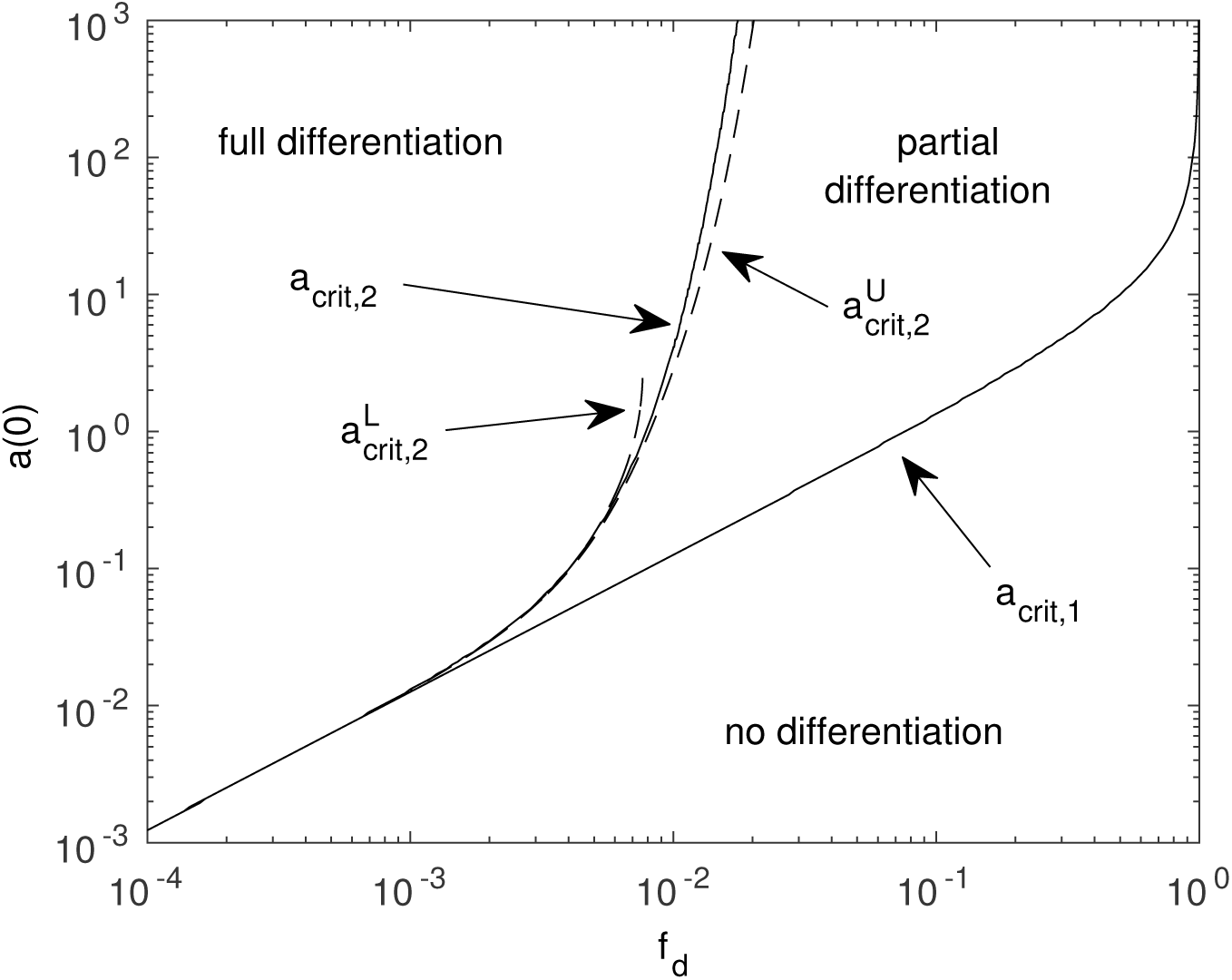
The critical values of *a*(0) shown against the threshold parameter *f*_*d*_ overlaid with the approximate values for *a*_crit,2_determined in Section 4.1.2, namely the upper estimate 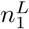 and lower estimate 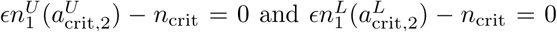 which are both shown as dashed linesHere, *f*_*p*_ = *f*_*d*_ + 4 *×* 10^-2^ and all other parameter values are given in Table 3.

A similar procedure can be performed with the lower bound on *f*_0_ to obtain a lower estimate 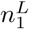 for the number of chondrocytes produced. Both these estimates are are in good agreement with the numerical solution at moderate amplitudes of *a*(0) (see Fig. 9(c)), and only diverge from the computed value at larger concentrations.

Finally, upper and lower estimates of the critical value *a*_crit,2_ correspond to the values of *a*(0) such that 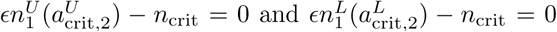, respectively. It is straightforward to find these values numerically in MATLAB, and the value of 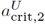 from the upper approximation (vertical dotted line in Fig. 9(c)) slightly underestimates the numerically obtained value of this parameter. In the example showncrit,21no approximation of 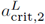 is possible since *n*^*L*^ does not reach the critical value *n*_crit_. The upper and lowerapproximations of *a*_crit,2_ are shown against the differentiation threshold parameter *f*_*d*_ in Fig. 10. Both are excellent approximations to the numerical solution for moderate amplitudes of *a*(0), and 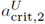 remains closeto the numerical value even for larger amplitudes, with a maximum relative error in the upper bound of15.5% for the largest amplitude of *a*(0) shown here. As in Fig. 9(c), no lower approximation 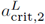 existsbeyond a moderate value of *f*_*d*_ as the lower bound on *n*_1_ never reaches *n*_crit_, even for large values of *a*(0).

To summarise, this analysis replicates the behaviour seen in the numerical simulations from Section 2.1.2 where full differentiation of a MSC population is achieved if sufficient exogenous TGF-*β* is added. The key insight that this analysis yields is an approximate expression for the concentration above which full differentiation takes place, as well as revealing the dependence of this value on model parameters. This estimate has been calculated by determining how many cells are produced by an initial concentration of exogenous TGF-*β* as described in Equation (64).

## 5. Hybrid strategy: add exogenous TGF-*β* to a co-culture

A final chondrogenesis strategy combines the strategies from section 3 and 4, as in the experimental study of [12] where a mixed cell population of 25% chondrocytes and 75% MSCs was cultured with various doses of TGF-*β* up to 10 ng/mL. The intent of this combined strategy is that is will be more cost-effective and robust than relying on driving chondrogenesis by either the use of co-cultures, or the addition of exogenous TGF-*β* alone; where growth factors are expensive and harvesting of chondrocytes is difficult it is likely that a hybrid approach will offer numerous advantages.

This situation is straightforward to implement in our model. We first obtain the fast kinetics values for all the TGF-*β* related quantities for a given initial co-culture population (24)–(28). Here, we assume that the number of seeded chondrocytes is less than *n*_crit_, so the co-culture does not completely differentiate without any extra stimuli. The effect of adding extra exogenous TGF-*β* to this system is then examined by a using fast kinetics values for *c*, *b*, *f*, *g*, *m* and *n* as initial conditions in the full numerical simulation, with the initial value for *a*, *a*(0) = *a*_FK_ + *a*_exogenous_, perturbed from its fast kinetics value by the concentration of the exogenously added TGF-*β*.

The results of some example simulations are shown in Fig. 11, where this hybrid strategy is compared to the strategy of adding exogenous TGF-*β* to a population of pure MSCs. As shown in Fig. 11(a), the value above which any extra chondrocytes are produced *a*_crit,1_ is almost unchanged by seeding of the 3% chondrocytes, but the value for *a*_crit,2_ is decreased by around a factor of 2. This indicates that combining these two strategies does indeed dramatically decrease the amount of TFG-*β* required to induce the full differentiation of the population.

**Figure 11:**
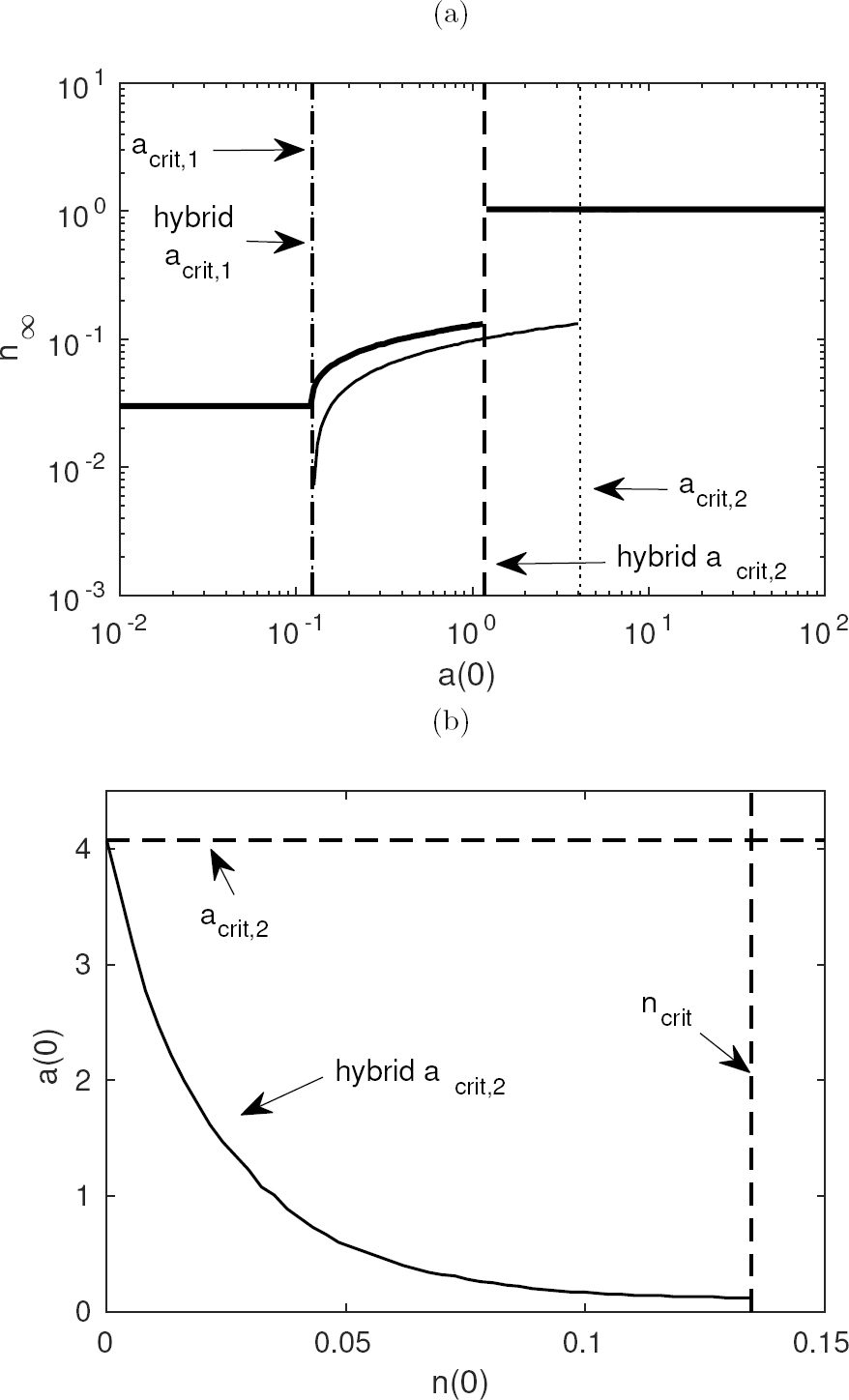
Simulations for the case of a combination strategy of both adding chondrocytes and exogenous TGF-*β*. (a) Showing the chondrocyte yield against initial concentration of TGF-*β* for two initial chondrocyte densities; the thin solid lines are for *n*(0) = 0 and the thick solid lines are for *n*(0) = 0.03, with the critical values for these two cases shown as dotted and dashed lines, respectively. Here the value of “hybrid *a*_crit,2_” for the case where some chondrocytes were in the initial cell population is significantly lower than for the case without any chondrocytes initially. (b) The dependence of *a*_crit,2_ for a range of initial chondrocyte densities *n*(0) for a fixed value of *f*_*d*_ = 10^-2^. The value of the hybrid *a*_crit,2_ decreases markedly as the number of seeded chondrocytes is increased. Full differentiation occurs if *a*(0) and *n*(0) are chosen so that they lie above the solid curve. Here, *f_p_* = 5 *×* 10^-2^ and all other parameter values are given in Table 3.

Some further results are shown in Fig. 11(b) indicating how the cut-off value for full differentiation, *a*_crit,2_, changes as the proportion of initially seeded chondrocytes and the concentration of exogenous TGF-*β* is increased. For a fixed value of the differentiation threshold parameter *f*_*d*_ = 10^-2^ the amount of exogenous TGF-*β* required to induce complete differentiation drops by more than an order of magnitude for a co-culture seeded with 10% chondrocytes, as compared to the case with purely MSCs. This prediction seems reasonable in light of the modelling of the individual strategies in isolation in Sections 3 and 4, and could result in cost-savings through the use of fewer chondrocytes and lower of TGF-*β* concentrations. Experimentally testing of this mixed strategy would be an interesting direction for future work.

## 6. Discussion and conclusions

We have developed a model to examine the differentiation of a population of MSCs into chondrocytes in response to TGF-*β*. There are two key points of novelty in this model. Firstly, the concentrations of the various forms of TGF-*β* are modelled explicitly, so that it is possible to track the storage of TGF-*β* in the ECM in a latent form for later activation. Secondly, the key aspects of the receptor kinetics involved are captured in a simple way, namely by the use of step functions to trigger differentiation or proliferation when the TGF-*β* bound to an MSC exceeds certain thresholds; this obviates the need to model the TGF-*β* signalling pathway explicitly. As discussed in Section 2, the use of step functions is standard practice in models of this type and has the advantage that the resulting model is amenable to detailed mathematical analysis. Possible issues with our model arise from the large number of parameters required to describe the complex biochemistry of the system. Although the values of these parameters were estimated from the literature, there is naturally a degree of uncertainty about the accuracy of these estimates. Similarly, the assumption that the cell populations and chemical concentrations are well-mixed means that any spatial effects (diffusion of chemicals for instance) are not considered here.

Two commonly used experimental chondrogenesis strategies were simulated. Our analysis of chondrogenesis driven by co-culturing MSCs and chondrocytes in Section 3 revealed that there is a critical initial density of chondrocytes *n*_crit_, given in (31), required to induce a population of MSCs to completely differentiate. The implication of this result is that if the initial cell density is above this critical value then the long term cell population will consist only of chondrocytes, but if the initial density is below the critical value then no extra chondrocytes will be produced and the long term cell population will be identical to the initial population. Thus, the optimal strategy suggested by the model is to add an initial density of chondrocytes which just exceeds the critical value; there is no real benefit to adding more chondrocytes that is sufficient to trigger chondrogenesis in terms of the ultimate yield or time taken to fully differentiate the MSCs, and so harvesting more chondrocytes than required to achieve this is unnecessary. The existence of such a critical value is broadly in line with behaviour seen in co-culture experiments. Under the conditions used in the *in vitro* study [29] the value of *n*_crit_ is between 5% and 25%, which is consistent with the values in Fig. 5(b). This suggests that the parameter values in Table 2 are not unrealistic and that a threshold parameter fordifferentiation *f*_*d*_ in the range 10^-3^–10^-2^ corresponds well with the results of *in vitro* studies.

Similarly, our analysis of the strategy of adding exogenous TGF-*β* to a population of MSCs revealed that in this excitable system the final yield of chondrocytes is dependent on the initial concentration of TGF-*β*, and that a population of MSCs will be completely differentiated if sufficient exogenous TGF-*β* is added. No cells are produced if the initial concentration is below a cut-off value *a*_crit,1_, and an analytic estimate of this value (see Equation (55)) was obtained. The entire MSC population can be driven to differentiation if the initial concentration exceeds a threshold value *a*_crit,2_, and a procedure to estimate this cut-off value, revealing its dependence on various modelling parameters, is given in Section 4.1.2. Our estimate of this second critical concentration, which is of particular interest in the context of tissue engineering, is slightly lower than the 10 ng/mL TGF-*β* conventionally added to *in vitro* experiments, suggesting that a similar outcome could be achieved experimentally with a lower concentration than is currently used. It would be interesting to test this experimentally, but this is beyond the scope of the present study.

Future extensions to this model could include an investigation of the effect of mechanical loading, for instance by using a piston to apply a load to cells within a hydrogel construct, as this kind of stimulation has been demonstrated to promote chondrogenesis [8] and has been proposed as a route to TGF-*β* activation [10]. Mechanical activation of this kind could be added to model of this paper by, for instance, replacing the chemical activation term in Equations (4) and (5) with a mechanotransductive term that depends on the local stress, meaning that TGF-*β* stored in the ECM could be activated and made available to cells in response to an applied mechanical load. This would necessarily require the extension of the well-mixed model of this paper, to one that includes spatial effects. A model which includes spatially varying cell densities and chemical concentrations would also allow other interesting biological features of the system to be investigated. For instance, this might include capturing the competition betweeen the diffusion of TGF-*β* and its rapid decay, which may mean that, for instance, TGF-*β* added to one side of a culture medium may not evenly spread throughout the medium, possibly resulting in isolated regions of the medium where noTGF-*β* is present. It would also allow consideratioin of cell movement by random motility and chemotaxis, both of which are likely be important in the context of cartilage tissue engineering to achieve a zonated cell distribution characteristic of cartilage, similar to that shown in Figure 1.

Since the synthesis of ECM components during chondrogenesis will dynamically change the bulk mechanical properties of extracellular environment [2, 30], further work is needed to couple the mechanical loading model with a model for the production of ECM components, in particular collagen type II and glycosaminoglycan. These extensions demand consideration of spatial effects, rather than the well mixed approach used in this paper, and developing such an approach will allow examination of further experimental questions of interest. For instance, in the context of tissue engineering it is necessary to understand the balance between the diffusion of exogenously added TGF-*β* through a medium and the rate at which it binds to that medium. Spatial dependence would also enable determination of the kind of stimuli required to drive some initial population of cells into a zonated final distribution of chondrocytes, as in natural articular cartilage.

This model elucidates a key component of the mechanism behind chondrogenesis and has revealed the excitable nature of this system. This is of particular interest in the context of tissue engineering and has great potential to refine experimental practice by predicting the concentration of TGF-*β*, or the number of chondrocytes, required to achieve full differentiation of a population of MSCs. This could, for instance, to cost savings via a reduction in the amount of TGF-*β* being added to a culture medium, or by harvesting fewer chondrocytes.

## Acknowledgements

The research leading to these results has received funding from the European Union Seventh Framework Programme (FP7/2007-2013) under grant agreement no. 309962 (HydroZONES). We thank Alexa Buβ from UK W urzburg for performing the chondrogenesis experiments shown in Fig. 2.

